# Variation in thermal sensitivity of diapause development among individuals and over time predicts life history timing in a univoltine insect

**DOI:** 10.1101/2023.05.31.543112

**Authors:** Jantina Toxopeus, Edwina J. Dowle, Lalitya Andaloori, Gregory J. Ragland

## Abstract

Physiological time is important for understanding the development and seasonal timing of ectothermic animals, but has largely been applied to developmental processes that occur during spring and summer such as morphogenesis. There is a substantial knowledge gap in the relationship between temperature and development during winter, a season that is increasingly impacted by climate change. Most temperate insects overwinter in diapause, a developmental process with little obvious morphological change. We used principles from the physiological time literature to measure and model the thermal sensitivity of diapause development rate in the apple maggot fly *Rhagoletis pomonella*, a univoltine fly whose diapause duration varies substantially within and among populations. We show that diapause duration can be predicted by modeling a relationship between temperature and development rate that is shifted towards lower temperatures compared to typical models of morphogenic, non-diapause development. However, incorporating interindividual variation and ontogenetic variation in the temperature-to-development-rate relationship was critical for accurately predicting fly emergence, as diapause development proceeded more quickly at high temperatures later in diapause. We conclude that the conceptual framework may be flexibly applied to other insects and discuss possible mechanisms of diapause timers and implications for phenology with warming winters.

## Introduction

Physiological time is an important concept for understanding the development and seasonal timing of ectothermic organisms (Reáumur 1735; Taylor 1981; Zhao et al. 2013; Buckley 2022). The cellular and physiological processes that underlie morphogenesis (development of an organism’s body size, shape and structure) are temperature-dependent (Kipyatkov and Lopatina 2010; Shi et al. 2011; Damos and Savopoulou-Soultani 2012). The vast majority of animals on our planet are ectothermic, so the chronological time (i.e., number of hours/days) required to reach a developmental landmark depends on environmental temperature. Most natural environments exhibit substantial temperature variation; chronological time is thus a poor metric of life history timing (van Straalen 1983; Trudgill et al. 2005; Rebaudo and Rabhi 2018; but see Cayton et al. 2015). Rather, ectothermic development is usually described in physiological time, which relies on quantifying the relationship between development rate (governed by physiological processes) and environmental temperature, and is often expressed in units of degree days (Taylor 1981; van Straalen 1983). Environmental temperature variability, duration of temperature exposure, and ontogeny (e.g., development stage) can complicate the calculation of physiological time (Kingsolver and Woods 2016; Sinclair et al. 2016; Buckley 2022), but physiological time models have been well-characterized in insects and can facilitate predictions of population dynamics during the growing season – i.e., during spring, summer, and fall (Powell and Bentz 2009; Régnière et al. 2012; Sridhar and Reddy 2013; Scranton and Amarasekare 2017). However, we have comparatively little insight into whether and how the same concept of physiological time applies to developmental processes that occur during winter. The relative dearth of information on the relationship between temperature and development during winter represents a substantial knowledge gap, especially if we want to understand the consequences of climate warming on insect phenology (Marshall et al. 2020; Buckley 2022).

### Diapause, quiescence, and their known relationships with temperature

Most temperate insects overwinter in diapause (Fig. 1 – green and yellow), a dormant, stress-tolerant state that delays growth, morphogenesis, and reproduction until conditions become more favorable (permissive) for these processes in spring or summer (Tauber and Tauber 1976; Hand et al. 2016; Wilsterman et al. 2021). Many insects spend the bulk of their life in diapause, either the majority of a calendar year (Fig. 1A) or sometimes multiple years (Tauber et al. 1986; Hanski 1988; Hahn and Denlinger 2011; Moraiti et al. 2014; Dupuis et al. 2016). If diapause ends during winter, the insect often enters another form of dormancy called ‘quiescence’ – a lack of morphogenesis due to non-permissive environmental conditions (Fig. 1B – dark blue; Jenkins et al. 2001; Hodek 2002; Koštál 2006). Unlike diapause, where dormancy is maintained during transient or early exposure to permissive environments, quiescent individuals are not recalcitrant and will immediately resume post-diapause processes such as growth and morphogenesis when conditions become permissive. Diapause and post-diapause quiescence can synchronize individuals to have similar life history timing, facilitating mate finding and supporting reproductive output (Tauber and Tauber 1976; Hand et al. 2016; Wilsterman et al. 2021). Given the prevalence of diapause in overwintering insects, any attempts to apply physiological time to overwintering development must consider this important process.

**Figure 1.**
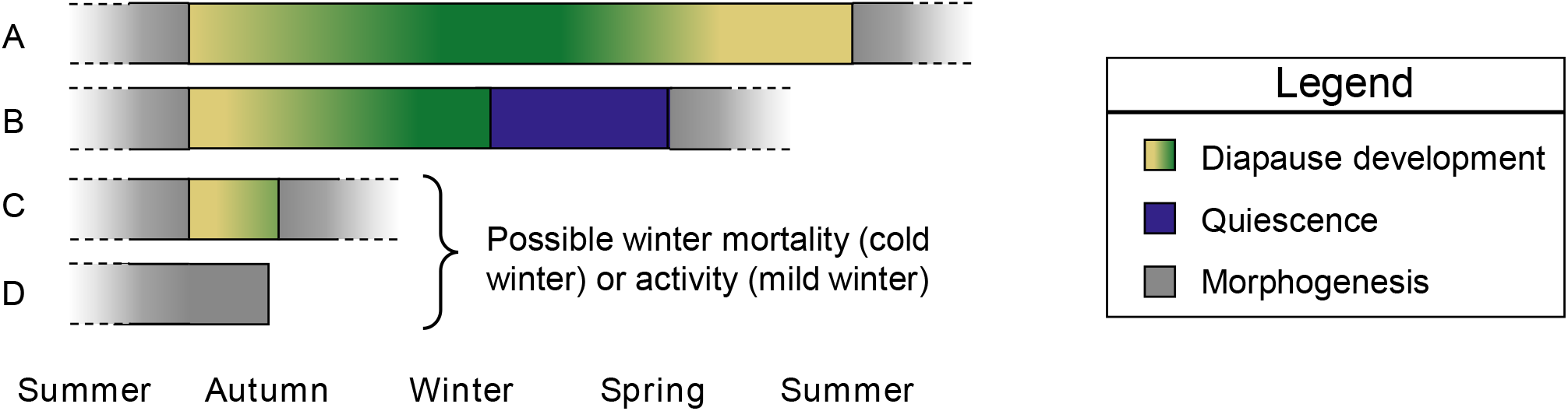
Example durations of diapause for insects that typically enter diapause in late summer or early autumn. **(A)** Some insects remain in diapause under warm (yellow) and cool (green) conditions for most of the year. **(B)** Many insects complete diapause during winter when it is cool (green), and enter a post-diapause dormancy called quiescence (blue), resuming post-diapause development (morphogenesis; gray) immediately when temperatures become permissive in the spring or early summer. **(C)** Warm summer or autumn conditions can cause individuals to complete diapause rapidly, referred to as ‘weak diapause’ or ‘shallow diapause’ in some species. **(D)** Non-diapause individuals never enter diapause and may complete morphogenesis prior to winter.

To expand the concept of physiological time to diapause, we must first establish that diapause is indeed a form of development. In contrast to growth and morphogenesis (morphological change; Fig. 1 – gray) during warmer or more permissive periods of the year, there is often no obvious morphogenesis during diapause (but see Shingleton et al. 2003), leading some authors to prefer alternatives to the term ‘diapause development’ (Hodek 1996). However, diapause is clearly a dynamic developmental process, where ‘development’ may primarily consist of progressive or cyclic changes at the cellular or molecular level (Andrewartha 1952; Hodek 1996; Koštál 2006), including differential expression of often thousands of genes over time (Koštál et al. 2017; Dowle et al. 2020; Pruisscher et al. 2022). Indeed, diapause can have distinct phases (e.g., initiation, maintenance, termination; cf. Koštál 2006), each of which must complete before post-diapause processes such as morphogenesis can resume. However, the mechanisms that determine the duration of each diapause phase (i.e., how the insect “counts” time during diapause) is still an area of active study (Hand et al. 2016).

Temperature has some well-described effects on diapause, but few studies have systematically investigated thermal sensitivity of diapause development. Chilling at low temperatures appears to be required for completion of the diapause program in many temperate species (Denlinger 2002), although there are many exceptions to this ‘rule’ (Hodek 2002; Zhu et al. 2009; Chen et al. 2014). In addition, warm temperatures can cause some insects to complete diapause relatively quickly (Fig. 1C) or never enter diapause (Fig. 1D; Denlinger 2002; Koštál 2006; Dambroski and Feder 2007; Toxopeus et al. 2021; Calvert et al. 2022). A common experimental approach manipulates chilling or winter durations to find the chilling threshold time required for diapause termination (e.g., Feder et al. 1997; Sgolastra et al. 2010; Higaki and Toyama 2012; Moraiti et al. 2014). Few studies (e.g., Lehmann et al. 2017) have expanded beyond these simple experimental designs to obtain insight into the underlying thermal sensitivity of the diapause development process, and the potential impact of warming autumn and winter conditions.

### Physiological time framework for morphogenesis

Models of physiological time based on the thermal sensitivity of morphogenesis are well-described and can inform our approach to modelling diapause development. Nonlinear, hump-shaped functions relating environmental temperature to morphogenic development rate generally fit empirical data quite well (Kipyatkov and Lopatina 2010; Shi et al. 2011; Damos and Savopoulou-Soultani 2012; Kingsolver and Woods 2016; Sinclair et al. 2016). These models, a class of thermal performance curves, predict zero development below or above the low and high temperature limits, respectively. Between these lower and upper thermal limits, as temperature increases morphogenic development rate increases to its highest value at the thermal maximum (c. 20 – 40 °C for many insects; Taylor 1981), and then sharply decreases back to a rate of zero as temperatures become too warm. Simpler slope plus intercept (straight line) models may also reasonably approximate the relationship between morphogenic development rate and temperature between the lower thermal limit and the thermal maximum (Trudgill et al. 2005; Kipyatkov and Lopatina 2010). The time to complete a developmental process at a particular temperature is the inverse of the development rate at that temperature (Kipyatkov and Lopatina 2010; Damos and Savopoulou-Soultani 2012), so development duration is shortest at the thermal maximum. We note that often authors refer to the thermal maximum as a thermal optimum, but we refrain from using “optimum” because maximal performance (e.g., fastest development) may not maximize fitness (Roff 1980).

When modelling morphogenic development rate, it is important to account for variation in the thermal sensitivity of development rate among individuals and over time. For example, genetic variation within populations can explain a substantial portion of interindividual variation in development duration that is often observed even under highly controlled laboratory conditions (Kingsolver et al. 2004). In addition, different life history stages (earlier vs. later in ontogeny) can exhibit different relationships between temperature and development rate that may be incorporated into predictive models (Yurk and Powell 2010; Lopatina et al. 2014; Sinclair et al. 2016; Kutcherov 2020). As with any thermal performance curve, the thermal maximum, the maximal performance value, or curve breadth may vary among individuals and across ontogeny. (Sinclair et al. 2012). Thermal generalists have a wide thermal performance curve (e.g., can develop across a broad range of temperatures), while thermal specialists have a narrow thermal performance curve, but often have a higher maximal performance at the thermal maximum (Sinclair et al. 2012).

### Proposed physiological time framework for diapause

We propose that diapause duration can be modelled using a diapause timer (cf. Hodek 2002) that is based on the principles of physiological time. To do this, we can use a thermal performance curve of diapause development rate. Given the well-known importance of chilling for diapause (described above), these diapause curves likely have very different shapes compared to functions describing morphogenic (non-diapause) development with maximum rates at high temperatures (cf. Hilbert et al. 1985). To our knowledge, there are no existing empirical estimates of thermal performance curves for diapause development, so we start with simple, hypothetical models that cover the most plausible forms of the relationship (Fig. 2 insets). Diapause phenology experiments often involve incubating insects at two different, consecutive temperatures (chilling then warming; e.g., Rull et al. 2016; Lindestad et al. 2020; Toxopeus et al. 2021), so we present the predicted diapause and total development durations (in chronological time) associated with these simple thermal sensitivity experiments (Fig. 2), as expanded upon below.

**Figure 2.**
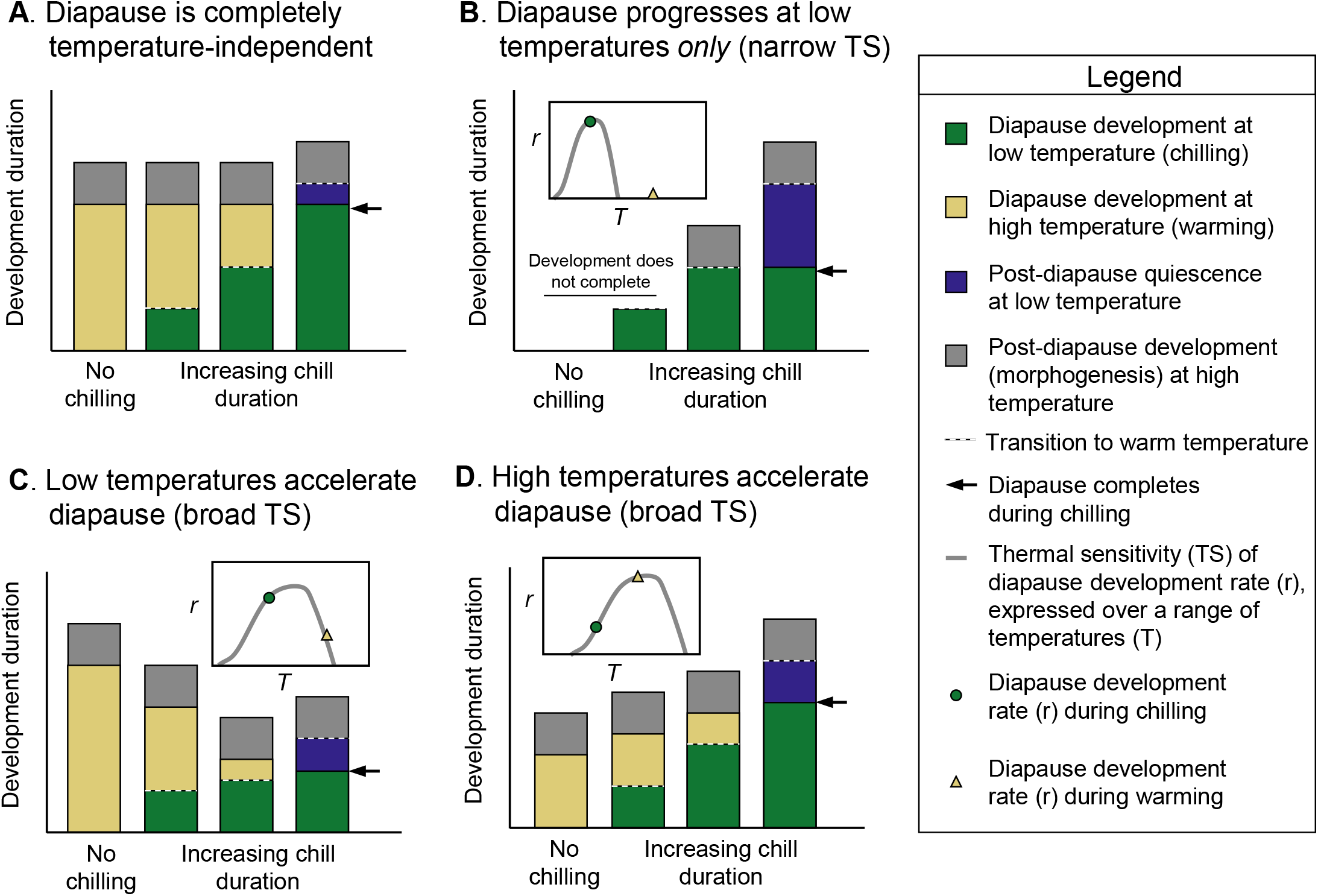
Hypothesized effect of a single (A) chronological or (B – D) physiological diapause timer on development duration under different combinations of chilling and warming. Transition from cold to warm temperature indicated by dashed line. Development duration (in chronological time) includes diapause development under cold (green) and warm (yellow) conditions, and post-diapause (e.g., morphogenic) development (gray) that can only occur under warm conditions. If individuals remain chilled after diapause completes (arrows), cold-induced post-diapause quiescence (dark blue) occurs until animals are returned to warm temperatures. Insets: relationship between temperature (T) and diapause development rate (r) at a low (chilling) and high (warming) temperatures based on an example thermal sensitivity (TS) of the diapause timer. The equations that explain the relationships between development rates, chill duration, and development duration are elaborated on in the Supplementary Information. Multiple diapause timers may exist (e.g., for different stages of diapause), which would result in more complex relationships between chilling duration and development duration (not shown).

Our models make several simplifying assumptions about the environmental sensitivity of diapause and about how an observer measures diapause duration. We do not make any assumptions about how diapause is induced, but do assume that temperature alone (and not photoperiod or other cues) influences the duration of diapause. This assumption applies reasonably well to many insects that complete diapause without additional environmental cues regardless of whether diapause was induced by a non-temperature cue (Tauber and Tauber 1976; Hodek 1996; Denlinger 2022). Directly observing the precise timing of the end of diapause is often not practical. This is because overt, morphological changes (morphogenesis) associated with resumption of post-diapause growth and development often occur well after hormonal and metabolic pulses that delineate the true physio-developmental transition (Hodek 1996, 2002; Denlinger 2002). Thus, we further assume that a measurement proxy for diapause duration also includes some amount of post-diapause development (morphogenesis; Fig. 2-gray), culminating in a clear phenotypic marker such as a life stage transition (e.g., adult emergence of a pupal diapausing insect) or a behavior (e.g., emergence from hibernacula). Finally, we assume that though diapause may progress at relatively cold temperatures, post-diapause development only proceeds at relatively warm temperatures. If diapause ends during low temperature exposure, quiescence occurs (Fig. 2 – dark blue), which delays post-diapause development because low temperatures impede morphogenesis (Jenkins et al. 2001; Hodek 2002; Koštál 2006).

Consider a naïve model in which diapause development progresses at a constant rate over time, independent of temperature. Under this chronological timer, diapause duration is approximately the same regardless of thermal environment (Fig. 2A) – some proportion of diapause development (and post-diapause development) occurs during chilling and the rest of diapause development occurs during warming. The only treatment that can increase overall development duration is chilling the insects for longer than their diapause duration (e.g., Fig. 2A; longest chill duration), which delays resumption of post-diapause development while insects are in chilling-induced quiescence. More nuanced chronological diapause timers are possible, e.g., chronological timers that are “started” by chilling or warming, but we focus here on the simplest model as a point of comparison to models that incorporate physiological time (Fig. 2B-D).

Realistically, we expect temperature to influence diapause development in some way. First, consider the classic “chill-dependent” diapause (or chilling threshold) model (cf. Dambroski and Feder 2007), where diapause can only progress at low temperatures. In this case, we predict that the physiological diapause timer has a fairly narrow thermal sensitivity, such that the rate of diapause development (*r*) is high at low temperatures, but zero at high temperatures (Fig. 2B inset). Individuals must be chilled for a sufficient time (chilling threshold) to complete diapause, and those that experience no or short chilling durations never complete 100% of the diapause program, and therefore cannot proceed to post-diapause development when warmed (Fig. 2B). Alternatively, the physiological diapause timer might have a broad thermal sensitivity, with a thermal maximum for diapause development rate at low (Fig. 2C inset) or high (Fig. 2D inset) temperatures. If low temperatures accelerate the diapause timer, diapause duration and chilling duration have an inverse relationship: long chilling durations result in short total development durations (Fig. 2C). Under this diapause timer, the longer the individual is in a cold environment, the greater the proportion of diapause completed prior to warming, so less chronological time is needed at warm temperatures to complete diapause. If high temperatures accelerate the diapause timer, we expect a positive correlation between chill duration and total development duration, with the shortest diapause duration in individuals that are only exposed to warm temperatures due to high rates of diapause development at these temperatures (Fig. 2D).

Just as thermal sensitivity may vary among individuals and over time during morphogenic development (Yurk and Powell 2010; Lopatina et al. 2014; Sinclair et al. 2016; Kutcherov 2020), diapause development performance curves may differ among individuals and over time. For example, early diapause (e.g., initiation, maintenance) may have a different thermal sensitivity than late diapause (e.g., termination), which we have not captured in our simple models (Fig. 2). However, each thermal sensitivity model in Figure 2 can be independently applied to a distinct stage of diapause rather than diapause as a whole, if needed. We will refer to changes in the thermal sensitivity of development rate over time (e.g., due to transitions between diapause stages) as changes or variation across ontogeny.

### Study organism and goals

In this study we fit these naïve and physiological time models (Fig. 2) to measurements of diapause duration across simple cold/warm temperature treatments in the univoltine apple maggot fly *Rhagoletis pomonella* (Diptera; Tephritidae). *Rhagoletis pomonella* diapause as pupae under the soil for most of the year (Fig. 1A), and their diapause duration is governed primarily by temperature rather than photoperiod (Prokopy 1968; Feder et al. 1997), making *R. pomonella* a good model to examine the impact of temperature on diapause development. In addition, intra/inter-population variation in diapause related to seasonality has been intensively studied in *R. pomonella* (Dambroski and Feder 2007; Lyons-Sobaski and Berlocher 2009; Powell et al. 2020; Calvert et al. 2022). Our study suggests that diapause duration can be modelled using the principles of physiological time, with interindividual and ontogenetic variation as key components.

## Materials and Methods

### Rhagoletis pomonella as a diapause model

*Rhagoletis pomonella* has recently evolved to be an agricultural pest, laying its eggs in fruits of apple trees (*Malus domesticus*) that were introduced to North America several hundred years ago. It natively infests hawthorn (*Crataegus* spp.) trees throughout North America, and both apple- and haw-infesting populations mate and oviposit in the late summer/early autumn (Dean and Chapman 1973). Larvae develop within fruit, burrow into the soil to pupate, and then most pupae enter a long diapause and spend up to 10 months in the soil, eclosing as adults the following summer (e.g., Fig. 1A; Dean and Chapman 1973). Differences in life history timing (e.g., between apple- and hawthorn-infesting *R. pomonella*; Bush 1969; Feder et al. 1993) are driven by differences in diapause duration (Powell et al. 2020).

There are three well-described, apparently discrete diapause phenotypic classes in *R. pomonella* that vary in diapause intensity, or recalcitrance to develop despite permissive temperatures (Dambroski and Feder 2007; Calvert et al. 2022). In the absence of low temperatures, a small proportion (<10 %) of *R. pomonella* pupae designated non-diapause (ND) avert diapause completely (e.g., Fig. 1D; Prokopy 1968; Dambroski and Feder 2007; Calvert et al. 2022). Most pupae enter a prolonged chill-dependent diapause (CD) lasting up to 9-10 months in the field (e.g., Fig. 1A; Feder et al. 1997; Dambroski and Feder 2007; Calvert et al. 2022). A small portion (< 20 %) of pupae enter a weak diapause (WD; cf. Toxopeus et al. 2021), also termed shallow diapause (Dambroski and Feder 2007). These WD pupae will terminate diapause after only a few weeks if held at warm temperatures (e.g., Fig. 1C). Under constant warm conditions, these three diapause intensity phenotypes are relatively discrete, with multimodal distributions of eclosion and even distinct allele frequencies (Feder et al. 1997; Dambroski and Feder 2007; Calvert et al. 2022). However, continuous variation in physiological processes may underlie these discrete traits (Calvert et al. 2022), consistent with a threshold quantitative genetic model (Roff 1996).

### Insect collection and diapause incidence

We collected hawthorn fruits (*Crataegus* spp.) infested with *R. pomonella* in Denver CO during August 2017 and 2018, brought them to the lab, and collected emerging larvae and pupae as previously described (Toxopeus et al. 2021). Briefly, we placed fruits in wire mesh baskets suspended over plastic trays in an environmentally-controlled room (c. 22°C). We collected newly formed pupae three times per week and transferred them to a Percival DR-36VL incubator (Percival Scientific, Perry IA) set to 21°C, 14:10 L:D and 80% R.H. for 10 days. Diapause *R. pomonella* pupae suppress their metabolic rate during these 10 days, after which metabolic rate stabilizes (Ragland et al. 2009). We defined 10 d post-pupariation as “time zero” in our experiments. Temperatures during this early time period of diapause preparation and initiation can influence diapause development rate (Dambroski and Feder 2007; Ragland et al. 2012), but this study focuses on the influence of temperature on individuals that have already successfully entered diapause.

To estimate the proportion of *R. pomonella* pupae in each diapause phenotypic class (ND, WD, CD), we measured CO_2_ production in a subset of 2018 pupae at 10 d and 45 d post-pupariation at 21°C using stop-flow respirometry as previously described (Ragland et al. 2009; Toxopeus et al. 2021). A small number (6 out of 200) with intermediate metabolic rates were excluded because they could not be unambiguously assigned to a diapause phenotype. Because we focused our analysis on diapause development, we excluded ND pupae from subsequent analyses, either based on metabolic rate, or very short (< 40 days; cf. Feder et al. 1997; Toxopeus et al. 2021) times to complete development to adulthood after chilling treatments (see below).

### Temperature treatments and development tracking

#### Experiment 1: Different durations of chilling

To determine whether diapause was governed by physiological timers, and whether diapause development was accelerated at low or high temperatures, we compared the relationship between chilling duration and diapause duration in *R. pomonella* to the predictions in Figure 2. At time zero (10 d post-pupariation), pupae were separated into groups of c. 100 individuals. One group was kept at 21°C for the duration of the experiment (no chilling). All other groups were transferred directly to simulated winter (chilling at 4 °C and c. 85% R.H) for one of several chilling durations ranging from 2 to 29 weeks. We also included one experimental group (see Supplementary Information) that was chilled at a warmer temperature (15°C) for 9 weeks. Following chilling, pupae were transferred back to 21°C. Each experimental group was checked three times per week at 21°C for up to 125 d post-chilling to determine when adults eclosed from the puparia, the marker that diapause and post-diapause development (morphogenesis) were complete. No obvious defects in adult morphology were noted in any treatment. For each treatment group, we calculated the mean development duration (including time at 4°C and 21°C), the coefficient of variation (CoV) of mean development duration, and the proportion of pupae that eclosed as adults. Summaries of development duration only included individuals that eclosed as adults.

Mean diapause duration and the incidence of quiescence in *R. pomonella* was determined in each temperature treatment group based on two assumptions. 1) At 4°C, pupal morphogenesis (i.e., post-diapause development) cannot occur (Reissig et al. 1979), but diapause development can occur, based on the observation that pupae can complete development following prolonged chilling (Feder et al. 1997; Powell et al. 2020; Toxopeus et al. 2021). 2) At 21°C, the duration of post-diapause pupal morphogenesis is relatively invariant among individuals, so the duration of total development is largely driven by the chronological time it takes to complete diapause (Feder et al. 1997; Powell et al. 2020). If post-chill development duration is approximately equal to the time to complete morphogenesis (c. 30 days), we can infer that the diapause process completed at 4°C and post-diapause quiescence may have occurred (e.g., longest chill duration bars in Fig. 2). However, if the post-chill development duration is greater than that required to complete morphogenesis, we can infer that only a portion of the diapause process was completed at 4°C and the remaining proportion of diapause development occurred at 21°C prior to post-diapause morphogenesis (e.g., all chill durations except the longest in Fig. 2C, 2D). In this latter case, cold-induced quiescence does not occur, so diapause duration (e.g., green + yellow bars in Fig. 2) is the difference between age at eclosion (e.g., total height of bars in Fig. 2) and the duration of morphogenesis (e.g., gray bars in Fig. 2). We did not record body mass for any individuals for this or other experiments, though previous work suggests a positive relationship between body mass and emergence timing (Ragland et al. 2012).

#### Experiment 2: Chilling for the same duration but at different ages

To detect ontogenetic shifts in the thermal sensitivity of diapause development, we compared development duration of *R. pomonella* pupae exposed to the same chilling durations but at different chronological ages. We included additional temperature treatments in which groups of c. 100 pupae were kept at 21°C for a warming period (2, 3, or 6 weeks beyond time zero) prior to chilling (4°C) for 2, 3, 6, or 9 weeks. Pupae were then transferred back to 21°C and eclosion was tracked as described above. We compared development duration in these treatments to pupae that were chilled at time zero (10 d post-pupariation) for 2, 3, 6, or 9 weeks. If chilling at different chronological ages resulted in a decrease or increase in total development duration, this would suggest that the thermal sensitivity of the diapause timer can vary with ontogeny, i.e., it can change over the course of diapause development.

### Diapause timer computational simulations

Initial inspection of our empirical data suggested the results could not be explained by one of the simple relationships between diapause development rate and temperature (Fig. 2; see Results for additional details). Indeed, due to variation in the proportion of individuals that eclosed (completed diapause and post-diapause development) following each temperature treatment in Experiment 1 and the presence of WD and CD diapause phenotypes, it was clear that we needed to account for interindividual and ontogenetic variation in thermal sensitivity of diapause development. To do this, we developed a simulation model that incorporated these factors. The full derivation of the model equations and an explanation of model assumptions (summarized in Table S1) are available in the Supplementary Information, and briefly described below. All simulations and analyses were conducted in R v4.1.0 (R Core Team 2022), and our code is available as described in the Data and Code Accessibility statement.

Based on estimated proportions of diapause intensity phenotypes (see Results), we simulated a population that included 150 WD and 850 CD individuals, each with its own diapause timer(s) that specified diapause development rate at 4°C and 21°C. As Hodek (2002) noted, diapause development represents a decline in diapause intensity over time. Thus, including these discrete WD and CD phenotypes in the models is effectively adding a discrete dimension of diapause development rate variation in addition to the continuous interindividual variation described below. We modelled one diapause timer for each WD individual, and two diapause timers (early and late diapause development) for each CD individual, accounting for ontogenetic variation in diapause development in CD individuals. Within each diapause phenotype (WD, CD), diapause development rates varied among simulated individuals. In addition, we incorporated some chilling-related mortality for both diapause phenotypes based on empirical observations. Total development duration (diapause and post-diapause development) was calculated from development rates (Eq’ns 1.1 – 2.4) for each simulated individual that was ‘exposed’ to 4°C (chilling) at time zero for between 0 and 400 days, followed by warming at 21°C, similar to Experiment 1 above.

Total development duration (diapause and post-diapause development) of each WD individual as a function of duration at 4°C was simulated with the following equations that included a single physiological diapause timer:

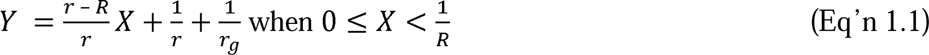

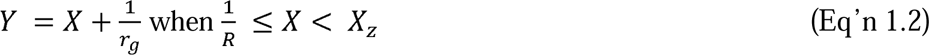

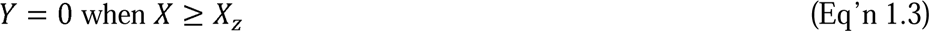

where *Y* is the duration (chronological time) to complete development since time zero, *X* is the duration of chilling (d), *R* is the rate of diapause development at 4°C (d^-1^; e.g., green circle in Fig. 2 insets), *r* is the rate of diapause development at 21°C (d^-1^; e.g., yellow triangle in Fig. 2 insets), *r_g_* is the rate of post-diapause development (morphogenesis) at 21°C (d^-1^), and *X_z_* is the duration of chilling (d) that causes mortality in WD individuals. The inverse of each rate gives the duration to complete that process (diapause or morphogenesis) at the respective temperature. When *Y* = 0 (e.g., Eq’n 1.3), that indicates that an individual was unable to complete development and adult eclosion never occurred, so development duration is effectively 0 d. We use *Y* = 0 rather than recording simulated individuals as dead because (as described below for CD individuals) some individuals fail to complete development because they are alive but temperatures are not permissive for development (e.g., no chilling and shortest chilling treatment in Fig. 2B).

To model total development duration (diapause and post-diapause development) of each CD individual, we included two physiological diapause timers – one each for early (*e*) and late (*l*) diapause development – in the following equations:

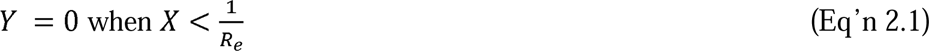

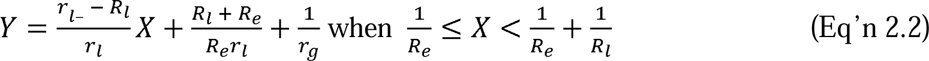

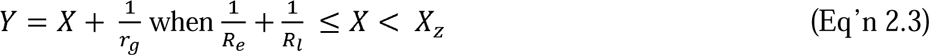

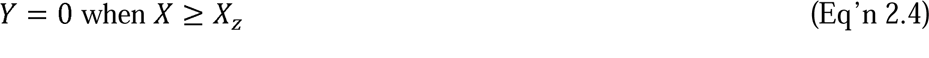

where *R_e_* is the rate of early diapause development at 4°C (d^-1^), *R_l_* is the rate of late diapause development at 4°C (d^-1^), *r_l_* is the rate of late diapause development at 21°C (d^-1^), *X_z_* is the duration of chilling (d) that causes mortality in CD individuals, and all other symbols are as described for the WD individuals. Early diapause must complete before late diapause can begin. We assumed that early diapause development did not proceed at warm temperatures (21°C), i.e., chilling was required for completion of early diapause (cf. Fig. 2B), and so individuals did not complete development and eclose (*Y* = 0) if chilling duration was insufficient (less than 1/*R_e_*) for individuals to complete early diapause (Eq’n 2.1).

In addition to discrete variation for the WD and CD phenotypes, we also modelled continuous variation in diapause development rate. The rates of diapause development for each of the simulated WD individuals (*R* and *r* in Eq’ns 1.1 – 1.2) and CD individuals (*r_l_*, *R_e_*, *R_l_* in Eq’ns 2.1 – 2.3) were randomly sampled from inverse Gaussian distributions of rates with the parameters described in Table S2. This approach proved most practical for accurately predicting the empirical data and is the focus of this study. We then calculated the same summary metrics (mean and CoV development duration, proportion eclosion) for our simulated population as for our empirical population for each chill duration used in Experiment 1, including only eclosed (*Y* > 0) individuals from summaries of development duration. We compared the fit of our simulation summary metrics to the empirical summary metrics by calculating the sum of squared residuals (SSR) of mean development duration and the χ^2^ of proportion eclosion between the two datasets. We used multiple iterations of these simulations to numerically solve for the parameters of our rate distributions (µ, λ in Table S2) that minimized the SSR and χ^2^ values.

While the assumptions in our final simulation model were justified by empirical data (see Supplementary Information), we tested the sensitivity of the model to these assumptions. To do this, we either modified or removed parameters from Eq’ns 1.1 – 2.4, ran our simulation, calculated the SSR and χ^2^ parameters as described above, and visually compared the fit of the simulation summary metrics to the empirical summary metrics. We also tested whether simpler models could produce appropriate patterns of proportion eclosion, and mean and CoV development duration, including: 1) only one chill-dependent diapause phenotype (Eq’ns 2.1 – 2.4) instead of distinct WD and CD phenotypes; and 2) two diapause phenotypes, but no ontogenetic variation in diapause development rates in either group. In addition, we attempted to fit a model that treated WD pupae as an extreme phenotype along a continuum of diapause development rates. The approaches to these alternative models and associated results are described in Supplementary Information. A subset of the most relevant alternative simulations is included in the main text.

## Results

The results detailed below broadly support the hypothesis that insect development during winter-like conditions can be explained by timers based on the principles of physiological time. The thermal sensitivity of these timers differ between diapause intensity phenotypes, among individuals, and across ontogeny. While there are multiple potential interpretations of the empirical data, below we focus on interpretations that were supported by our subsequent simulation models.

### Diapause timers vary among diapause intensity phenotypes

Based on the proportion of individuals that completed development under different chilling and warming treatments in Experiment 1, weak diapause (WD) and chill-dependent diapause (CD) phenotypes likely have different diapause timers. Consistent with a number of previous observations (Dambroski and Feder 2007; Toxopeus et al. 2021; Calvert et al. 2022), we confirmed the presence of ND (non-diapause), WD, and CD *R. pomonella* pupae reared under constant warm conditions using metabolic rate measurements (Fig. 3A). Under constant warmth, all ND flies completed their eclosion before WD flies began eclosing, and most CD flies remained in diapause under these conditions (did not eclose at all; Fig. S1, S2). CD pupae were most abundant (82%), followed by WD (15%), and ND (3%; Fig. 3B), the latter of which was excluded from our analyses (see Methods for details).

**Figure 3.**
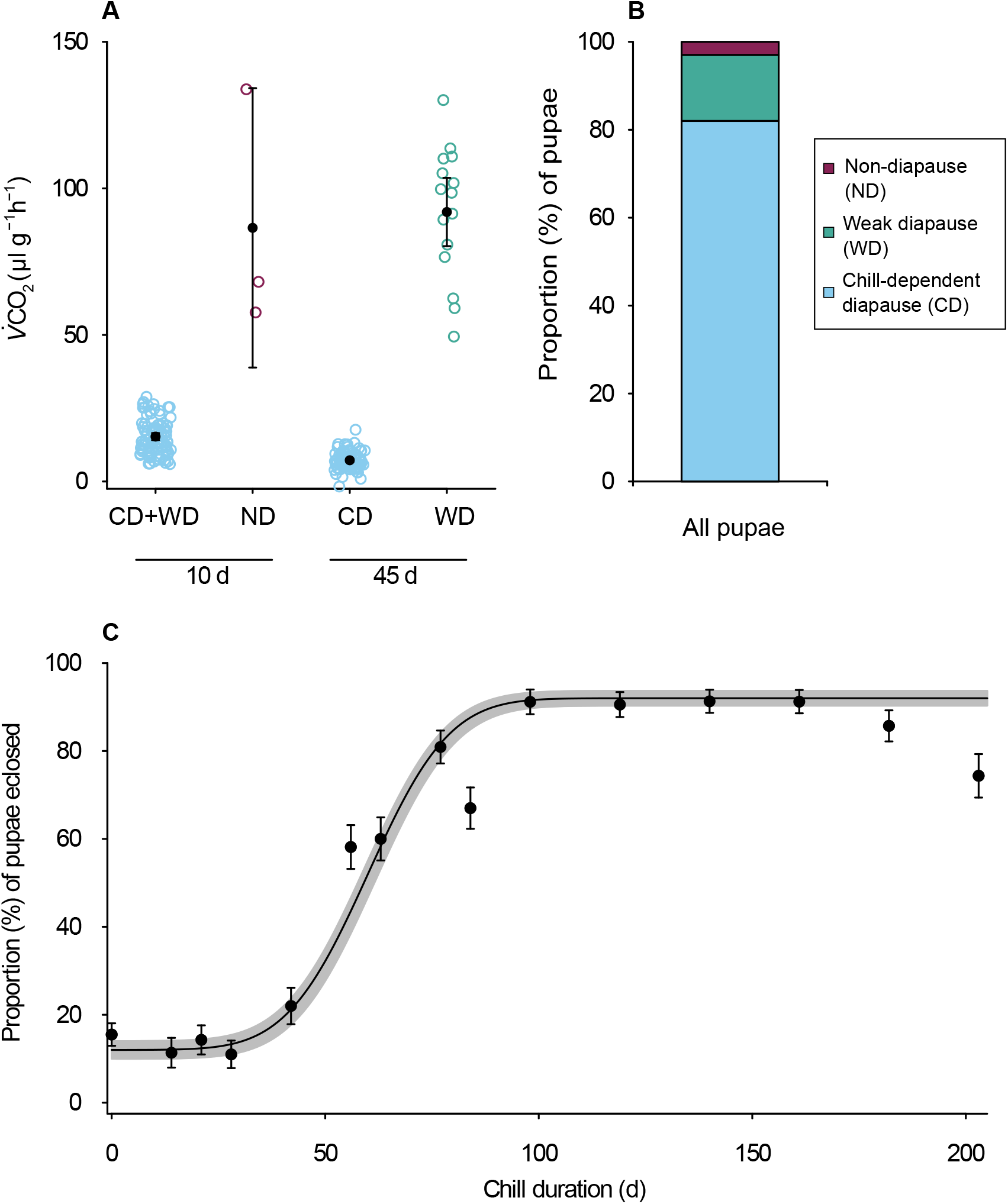
Diapause incidence and the effect of chilling on diapause completion in *Rhagoletis pomonella*. **(A)** Mass-specific metabolic rate after 10 or 45 days of constant warmth post-pupariation was used to determine diapause phenotype: non-diapause (ND), weak diapause (WD), and chill-dependent diapause (CD). Open circles, individual data points; filled circles, means ± 95% confidence intervals. Small error bars are obscured by symbols. **(B)** The proportion of ND, WD, and CD pupae inferred from the metabolic rate data in **(A)** and duration of total development (Fig. S2) in groups that were not metabolically-phenotyped. **(C)** Each point represents the proportion ± standard error of proportion of c. 100 pupae that completed diapause and post-diapause development (eclosed as adults) following exposure to chilling (4°C) for a specific duration at time zero (10 d post-pupariation). Black line is a cumulative distribution model with a mean ± standard deviation of 60 ± 15 d chilling. Gray shading indicates standard error of proportion of the model.

The two types of diapause timers in this population apparently included a rarer one that did not require chilling, and a common one that required moderate to prolonged chilling. As previously documented (Feder et al. 1997; Dambroski and Feder 2007; Toxopeus et al. 2021), some diapause *R. pomonella* pupae were able to complete diapause and eclose as adults following a range of temperature treatments, from no chilling (warming only) to prolonged chilling (>200 days) at 4°C (Fig. 3C). Approximately 15% of our population eclosed with little to no chilling (0 – 4 weeks at 4°C; Fig. 3C), which is similar to the proportion of WD pupae in our population (Fig. 3B). However, as chill duration increased, the proportion eclosion increased (Fig. 3C), suggesting that the remaining CD pupae required a threshold cold exposure (chilling) to successfully complete the diapause program. We confirmed that the majority of uneclosed pupae following short chill treatments appeared to be alive and undeveloped (Fig. S1), suggesting that they were unable to complete diapause, rather than dead, or developed but unable to leave the puparium. The small (c.10%) proportion of pupae that did not eclose after longer chill treatments died in their puparia (Fig. S1).

### Variation in diapause timers among individuals

The relationship between chill duration and proportion eclosion in Experiment 1 (Fig. 3C) also suggested that there was substantial interindividual variation in diapause timers among individuals within each diapause phenotype. This is best illustrated here in CD pupae, but our simulation models (see below) also confirm interindividual variation in WD diapause timers. If we assumed that 15% of eclosed *R. pomonella* are WD individuals (see Fig. 3B), any proportion eclosion above 15% indicated completion of development by CD individuals. The sigmoidal increase in proportion eclosion as chill duration increased can therefore be explained by continuous variation among CD individuals in rate of diapause development required to reach a chilling threshold at 4°C (Fig. 3C). Sixty days of chilling was required for 50% of flies to eclose, and more than 90 days of chilling resulted in maximum (91%) proportion eclosion (Fig. 3C).

### Diapause is governed by physiological timers

When *R. pomonella* were exposed to increasing chill durations in Experiment 1, the mean total development duration increased (Fig. 4), suggesting that the diapause timers were temperature-sensitive, as expected for physiological timers. However, low temperatures did not universally accelerate development (decrease mean development duration), as the classic chilling model (Fig. 2B) would predict. Indeed, at first glance, the positive relationship between chilling duration and mean total development duration (Fig. 4) is broadly consistent with a diapause timer that is accelerated by high temperatures and has a broad thermal sensitivity (Fig. 2D). However, as discussed above, the proportion of eclosed individuals varied with chill time due to the requirement for chilling in most (likely CD) individuals, which is partially consistent with the predictions of the classic, chilling only model (Fig. 2B). So, it seems that none of the simple models individually explain all of the observations, especially given that WD and CD pupae respond differently to chilling.

**Figure 4.**
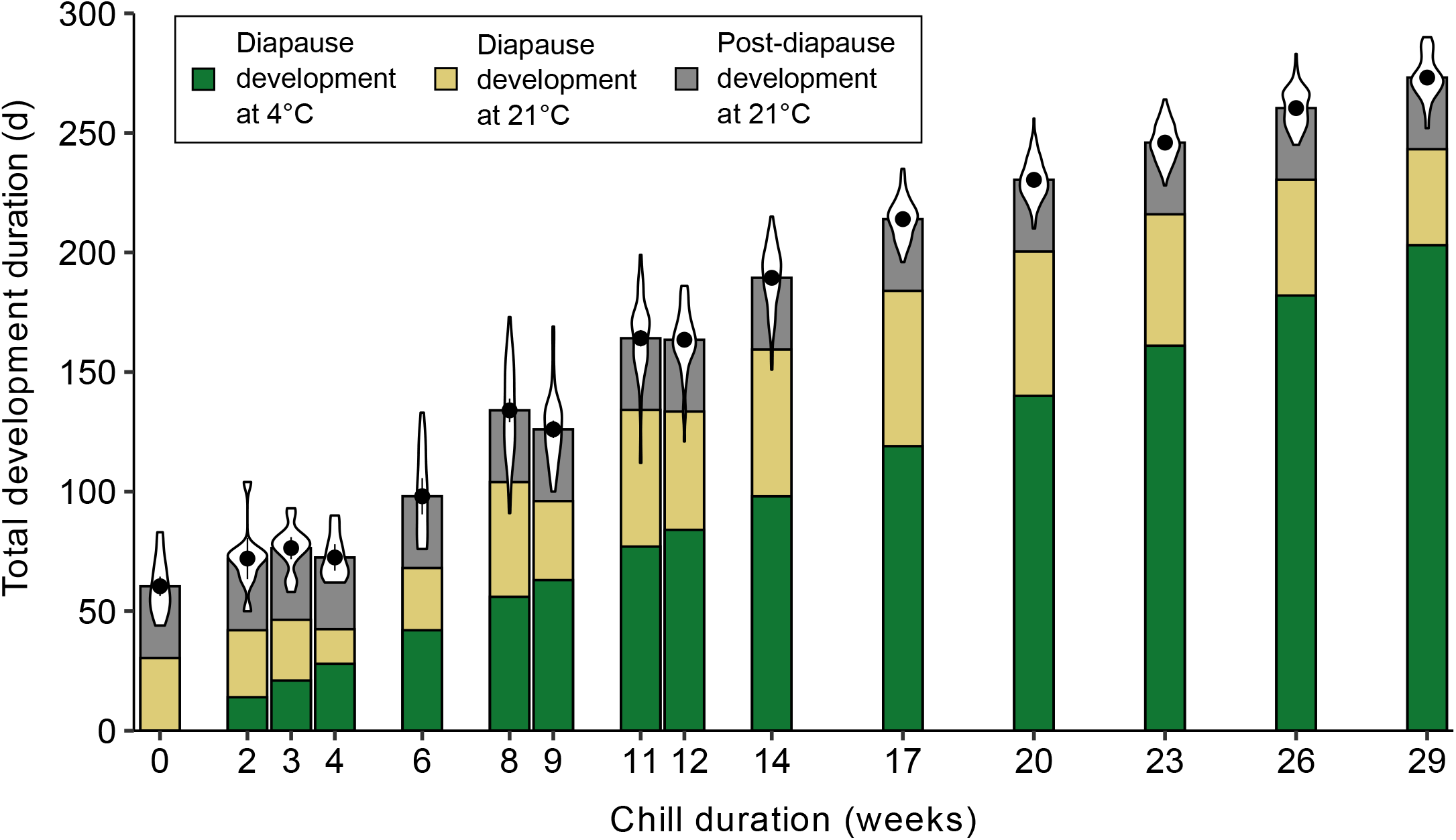
The effect of chilling on the mean duration of diapause and post-diapause development in *Rhagoletis pomonella*. Pupae were exposed to chilling for a specific duration at time zero (10 d post-pupariation) followed by warming. Each violin plot represents the distribution of durations to complete development (eclose as adults). Points are mean durations ± 95% confidence intervals. Small error bars are obscured by symbols. Bars below the violin plots show the mean duration of diapause development at 4°C (green – dictated by the chilling time) and 21°C (yellow - estimated). We assumed a mean value of 30 days for post-diapause development (morphogenesis; gray) at 21°C; gray bars are thus invariant across treatments.

### Diapause timers can have broad thermal sensitivity

Although chilling was required for completion of diapause in most of our population, diapause development appeared to have a broad thermal sensitivity (wide thermal breadth) for both WD and CD pupae. Under the assumption that all individuals completed post-diapause development in c. 30 days at 21°C (Powell et al. 2020), we reasoned that diapause could progress at 4°C and 21°C (green and yellow bars; Fig. 4) because the mean time to eclose following chilling (yellow + gray bars; Fig. 4) was always greater than the mean time to complete post-diapause morphogenesis at 21°C (gray bars; Fig. 4). We also found diapause could complete after “chilling” at the relatively warm temperature of 15°C (Fig. S3). None of our treatments in Experiment 1 (Fig. 4) included a clear period of cold-induced post-diapause quiescence (cf. longest chill treatments in Fig. 2). Indeed, data collected by Feder et al. (1997) suggested that more than 40 weeks of chilling are required for CD *R. pomonella* from hawthorn fruits to complete diapause at low temperatures and enter quiescence, which is longer than the maximum chilling duration (29 weeks) we used in our study.

### Thermal sensitivity of the diapause timer can change during ontogeny

While early chilling seemed to be required for completion of diapause in CD pupae (Fig. 3C), prolonged cold exposure slowed down diapause development (positive slope in Fig. 4). This suggested that CD pupae had at least two diapause timers: one that governed early diapause development and was accelerated by low temperatures (chilling), and another that governed late diapause development and was accelerated by high temperatures (warming).

Though a similar pattern is expected when comparing diapause to post-diapause development in most insects (Jenkins et al. 2001; Hodek 2002), we emphasize that this ontogenetic shift in thermal sensitivity of development rate clearly occurs during diapause in CD *R. pomonella* because our chill durations were too short to induce quiescence.

Indeed, when we exposed groups of *R. pomonella* pupae to identical cold treatments at different ages during Experiment 2, cold treatments later in development increased diapause duration (slowed development) compared to cold treatments initiated at time zero (Fig. 5). Because the age at chill exposure affected total development duration, this suggested that the rate of diapause development at low temperatures could change during diapause ontogeny. However, there were multiple factors at play that could influence mean development duration in these treatments (Fig. 5), including differences in the proportion of individuals that eclosed (Fig. S1) and the proportion of eclosed individuals that were CD vs. WD. Disentangling these effects empirically was challenging because it is only possible to metabolically phenotype CD and WD accurately when these individuals are held at warm temperatures for at least 45 days pre-chill (Dambroski and Feder 2007; Toxopeus et al. 2021; Calvert et al. 2022), which we could not do in all treatments. To address this issue, we developed simulation models to investigate whether interindividual and ontogenetic variation in diapause timer thermal sensitivity could explain the trends we observed in our empirical diapause development data.

**Figure 5.**
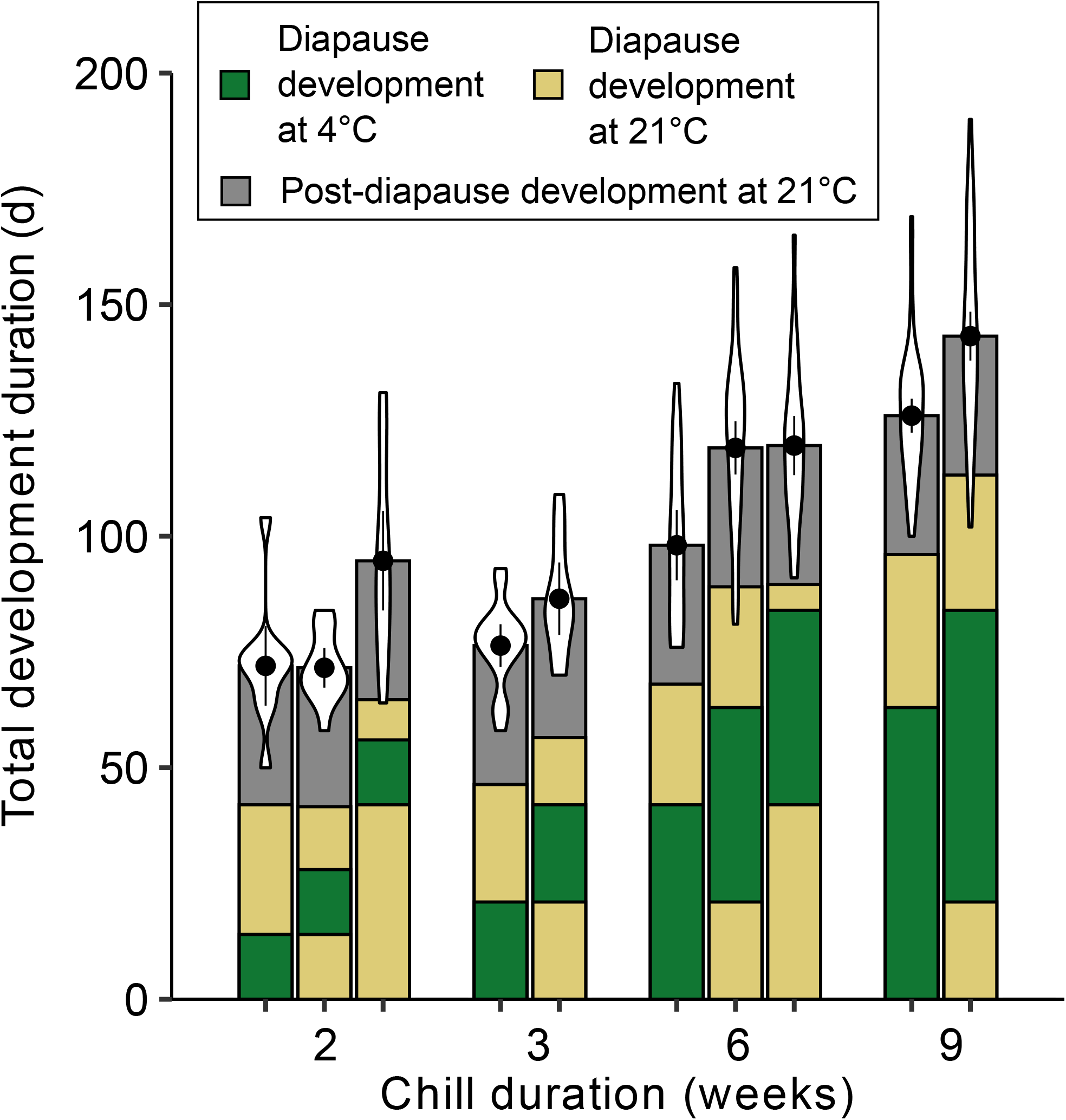
The effect of chilling at different ages on the mean duration of diapause and post-diapause development in *Rhagoletis pomonella*. Pupae were exposed to chilling for a specific duration at time zero (10 d post-pupariation), or 2, 3, or 6 weeks after time zero followed by warming. Each violin plot represents the distribution of times to complete from 10 d post-pupariation to adult eclosion. Points are mean durations ± 95% confidence intervals. Bars below the violin plots show the mean duration of diapause development at 4°C (green – dictated by chilling time) and 21°C (yellow - estimated), and assumed duration of morphogenesis at 21°C (gray).

### Accurate models must incorporate variation in diapause timers between diapause phenotypes, among individuals, and across ontogeny of diapause development

A simulation model incorporating interindividual and ontogenetic variation in diapause development rates, as well as differences between diapause phenotypes, provided an exceptional fit to our empirical observations. In the simulated population, each WD pupa had one diapause timer with a broad thermal sensitivity (Fig. 6A), while each CD pupa had two diapause timers: one for early diapause development (pre-threshold) progressing only at low temperatures, and one for late diapause development with a broad thermal sensitivity (Fig. 6B). The WD rate of diapause development was approximately twice as high at 21°C than 4°C (Fig. 6A), resulting in faster completion of diapause (shorter development durations) at 21°C (Fig. 6C). For simulated CD pupae, early diapause required 68.2 ± 16.4 d of chilling to complete (Fig. 6C), and the subsequent late diapause development rates were approximately four times higher at the warmer temperature (Fig. 6B), resulting in faster completion of late diapause at 21°C than 4°C (Fig. 6C). The overlap between thermal sensitivity of WD and late CD diapause development and that of post-diapause development (Reissig et al. 1979) allows completion of the diapause program at warm temperatures, followed immediately by post-diapause development. We note that our simulated diapause development rates were only modelled at two temperatures; more complex linear (e.g., polynomials) or nonlinear models would almost surely provide better fits over a broader temperature range (Lehmann et al., 2017; cf. Fig. 6A, B).

**Figure 6.**
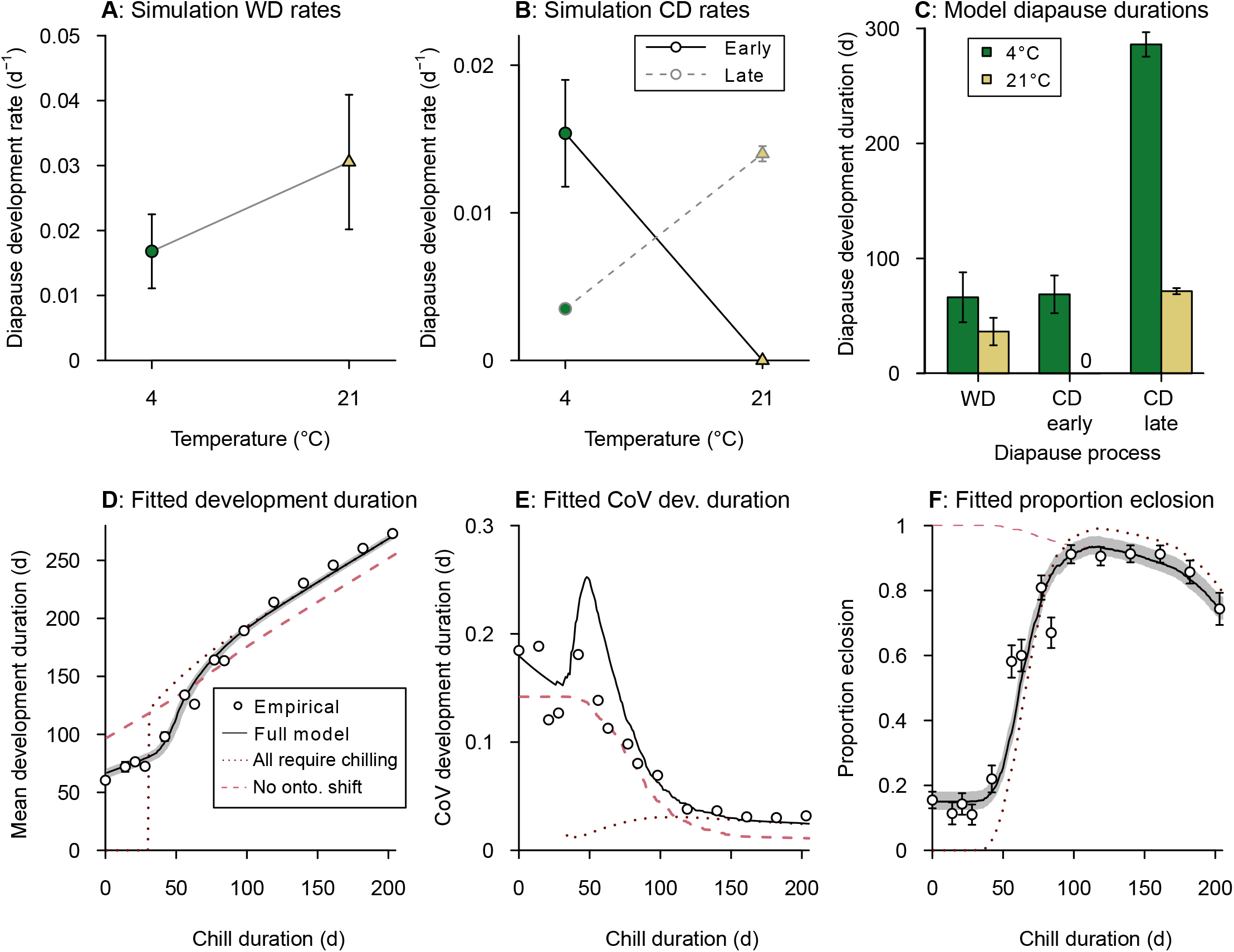
Summary of simulated diapause development in *Rhagoletis pomonella*. Mean and standard deviation of diapause development rates in a simulated population under cold (green) and warm (yellow) conditions for **(A)** 150 weak diapause (WD) pupae and **(B)** 850 chill-dependent diapause (CD) pupae during early and late diapause development. **(C)** Mean and standard deviation of diapause duration in simulated WD pupae and duration of early and late diapause in simulated CD pupae at 4°C or 21°C. 0 indicates failure to complete the developmental process. **(D-F)** Summary metrics of development duration and proportion eclosion phenotypes versus chill time for empirical data (symbols) and fitted simulation models (lines). The full simulation model (black line, Eq’ns 1.1 – 2.4) is compared to models that only included a single chill-dependent diapause phenotype (purple dotted line) or did not allow for ontogenetic shifts in thermal sensitivity of diapause development rates (pink dashed line). Gray shading indicates the 95% confidence interval of the simulation summary metrics in (D) and (F). Error bars represent standard error of the (D) mean or (F) proportion in empirical data. Small error bars are obscured by symbols. CoV, coefficient of variation of mean development duration.

The simulation model was sensitive to some, but not all, of our assumptions about interindividual variation in diapause development rate (Table S1). We modelled diapause development rates with inverse Gaussian distributions, but the final simulation model fit to empirical data did not differ substantially if we modelled rates with Gaussian distributions instead (Fig. S4). We also assumed that development rates within an individual were correlated across temperature (and time, in the case of CD pupae), but the final simulation model fit to empirical data was not sensitive to this assumption (Figs. S5, S6). Conversely, high interindividual variation in diapause development rate at the temperatures closer to the thermal maxima, and high interindividual variation in CD early diapause (see error bars in Fig. 6A, B) were both important for the final simulation model fit to empirical data (Figs. S5, S6).

Using the full model described in the methods, our simulated population (Fig. 6A, B) produced a close fit to the empirical data for mean development duration (Fig. 6D), variation in development duration (Fig. 6E), and proportion eclosion (Fig. 6F) following different chilling durations. Our full simulation model captured the prominent discontinuity/transition in development duration (Fig. 6D) and proportion eclosion (Fig. 6F) at c. 60 days chilling caused by a transition in high proportion of WD to high proportion of CD phenotypes among eclosed individuals. It likewise captured the decline in proportion eclosion at long chill times (Fig. 6F) driven by overwinter mortality. The simulated variation in development duration (Fig. 6E) was over-estimated for moderate chill treatments (6 – 14 weeks). Attempts to modify the distributions of diapause development rates to improve the fit for development duration metrics resulted in poor fit for proportion eclosion or CoV metrics (Figs. S7-9).

Alternative patterns of variation in diapause thermal sensitivity also did not improve fit. If we modelled two discrete diapause phenotypes, but did not allow for ontogenetic variation in rates for either group, we could not achieve the discontinuity in mean development duration associated with moderate chill lengths (Fig. 6D). If we only modelled a single chill-dependent diapause phenotype with an inverse Gaussian distribution of development rates, we could not recreate the pattern of c. 15% proportion eclosion after little to no chilling (Fig. 6F). We were able to improve the fit if we treated WD pupae as an extreme phenotype along a continuum of diapause development rates, but this model still did not accurately predict eclosion with little or no chilling (Fig. S10). Further efforts devoted to describing and modelling an unknown, continuous distribution of diapause development rates could lead to more generalizable models in this or other insect species. However, these efforts would not change the core principles of our model: diapause is a temperature-sensitive process that varies among individuals, across diapause phenotypes, and – in the case of CD pupae – across ontogeny.

## Discussion

Our study demonstrates that principles of physiological time that predict development duration during the growing season in ectotherms can also predict diapause development duration under various simulated winter conditions. Here we use data from *R. pomonella* to parameterize well-fitting models that incorporate similar principles. Our experimental design exposing each group of *R. pomonella* to two temperatures (chilling and warming) for different durations and at different ages facilitated detection of at least two distinct phases of diapause development, each with its own thermal sensitivity of diapause development rate. It likely would have been difficult to detect this ontogenetic shift in thermal sensitivity of diapause development in a classic thermal performance curve experiment, where multiple temperatures treatments were used but each group of insects was exposed to a single temperature (e.g., Irwin et al. 2001; Sgolastra et al. 2010; Xiao et al. 2013; Lehmann et al. 2017). Although we do not know the exact shape of the diapause development rate thermal performance curves in *R. pomonella*, our models of development rate at two temperatures are sufficient to explain most of the impacts of chilling and warming. Further work modelling the relationship over a broader temperature range may substantially improve predictions in variable and changing environments, and could perhaps extend to diapause occurring during warm seasons (Masaki 1980) as well.

### Extensibility of the model

The conceptual model of thermally sensitive diapause development with rate variation over time is consistent with observations in other insects. Several studies have shown that different overwintering temperatures impact diapause duration, with the fastest diapause development (shortest diapause duration) at temperatures between 0°C and 7°C (Irwin et al. 2001; Sgolastra et al. 2010; Xiao et al. 2013; Lehmann et al. 2017). Others provide evidence that the relationship between diapause development rate and temperature changes over time during diapause (e.g., Nomura and Ishikawa 2000; Xiao et al. 2013; reviewed extensively by Hodek 2002). The ‘early’ and ‘late’ phases we describe in *R. pomonella* could be consistent with a transition from the maintenance to the termination phase in a common, highly cited conceptual model of diapause development (Koštál 2006). Alternatively, ontogenetic variation in thermal sensitivity could be continuous – our experimental design could not test this possibility. The nature of ontogenetic variation in diapause development may not be settled, but we predict that changes in development rate during diapause are likely widespread, especially in univoltine species whose diapause continues across multiple seasons.

Similarly, we posit that parameterizing models to include interindividual variation may often provide more accurate and complete predictions of the distribution of diapause timing for an insect population. Many studies present evidence for pronounced interindividual variation in multiple aspects of insect diapause (e.g., Bradshaw and Lounibos 1977; Vinogradova 1986; Dingle and Mousseau 1994; Schmidt et al. 2005), including the thermal sensitivity of diapause duration (Nomura and Ishikawa 2000; Irwin et al. 2001; Masaki 2002; Chen et al. 2013; Lehmann et al., 2017). In this study we modelled continuous (e.g., variation within CD pupae) and discrete variation (WD vs. CD puape) in the thermal sensitivity of diapause development rate. Attempts to model a single diapause phenotypic class (i.e., with WD-like individuals having the fastest development rates along a continuum) provided a reasonable fit to some, though not all, of the data (Fig. S10). We also emphasize that we designed our experiments to test for simultaneous effects of interindividual ontogenetic variation, not strictly to predict field phenology. Thus, some details of our *R. pomonella* model may be idiosyncratic, but the conceptual framework is flexible, and we expect that it can be extended to other species. Ultimately, confronting models with data from the laboratory and the field across multiple species will be necessary to assess model generality.

### Implications for life history timing in changing environments

Increasing and more variable environmental temperatures across the entire year will likely affect life history timing. Warm pre-winter conditions can negatively impact some insects by causing aversion or early termination of diapause (Denlinger 2002; Koštál 2006; Toxopeus et al. 2021). In addition, warm temperatures can cause energy drain that leads to mortality (Sinclair 2015; Roberts et al. 2021; Nielsen et al. 2022). As autumn temperatures increase and winter warm spells become more frequent (Williams et al. 2015; Marshall et al. 2020), early diapause development in chill-dependent *R. pomonella* will likely slow down, resulting in an overall delay in the resumption of post-diapause morphogenesis in the following summer or fall, and a possible mismatch in adult eclosion and host fruit availability. Conversely, the predicted global increase in spring temperatures (Williams et al. 2015; Marshall et al. 2020) may shorten diapause duration in *R. pomonella* because this warming is likely to coincide with the later diapause development process that is accelerated at high temperatures. Thus, any models that attempt to predict insect phenology based on rates of post-diapause morphogenesis (e.g., Reissig et al. 1979) should take diapause duration into account, especially for species with a temperature-dependent diapause termination (Lehmann et al. 2017).

### How do insects ‘count’ the cold?

The physiological or cellular processes that determine diapause duration remain elusive (Hand et al. 2016), but some studies support a temperature-sensitive timer. For example, the time to complete chill-dependent diapause may depend on a particular process that is active at low temperatures. A single, key molecule accumulating or degrading over time might provide such a timer, for example, *aldose reductase* and *sorbitol dehydrogenase* in *Pyrrhocoris apterus* (Koštál et al. 2008), p26 protein in *Artemia franciscana* (King and MacRae 2012), or the metabolites alanine in *Pieris napi* (Lehmann et al. 2018) and sorbitol in *Bombyx mori* (Chino 1958). Alternatively, there could be a suite of genes or molecules regulating diapause duration. Gene expression changes extensively, sometimes transcriptome-wide during diapause (Robich and Denlinger 2005; Kim et al. 2006; Rinehart et al. 2007; Reynolds and Hand 2009; Emerson et al. 2010; Ragland et al. 2010; Urbanski et al. 2010; Kankare et al. 2016; Green and Kronforst 2019), including at low temperatures (Salminen et al. 2015; Koštál et al. 2017; Dowle et al. 2020; Pruisscher et al. 2022). Moreover, differentially expressed genes and genetic variants associated with diapause duration in *R. pomonella* are enriched in constituents of cell cycling and morphogenic pathways (Meyers et al. 2016; Dowle et al. 2020).

Our simulation and empirical datasets, and the transcriptomic studies cited above are both highly consistent with temperature-modulated developmental processes dictating the duration of diapause. We posit that this may be a general phenomenon. Diapause has evolved rapidly and repeatedly across the arthropod phylogeny (Ragland and Keep 2017; Denlinger 2022), suggesting that diapausing clades likely coopt existing developmental machinery from non-diapausing ancestors to initiate and regulate diapause development. This evolutionary hypothesis would also predict that models of physiological time would fit developmental data from both diapause and non-diapause development.

## Conclusions

Applying the principles of physiological time to overwintering *R. pomonella* revealed that diapause is temperature-sensitive, thermal sensitivity of diapause development rate varies among individuals and across ontogeny, and substantial portions of the diapause program progress more quickly at warm temperatures. Understanding the thermal sensitivity of overwintering development (even if that development occurs in a “dormant” state) is important for predicting how warming temperatures will impact the phenology of ectothermic animals. Additional work to test the generality of models and to elucidate the cellular and physiological basis for diapause timers is still needed, with implications for understanding how diapause has evolved.

## Acknowledgements

The authors would like to thank M. Calvert, M. Davenport, N. Sanaei, E. Rader, and M. Haigh for assistance with collecting and processing *R. pomonella* in the lab. We would also like to thank J. Kingsolver for comments on an earlier version of this manuscript, and V. Rudolf, S. Pincebourde, and two anonymous reviewers for comments on the submitted manuscript. This work was funded by National Science Foundation IOS 1700773 and DEB 1638951 grants to GJR.

## Statement of Authorship

JT, EJD, and GJR conceptualized the experiments. JT, EJD, and LA conducted the experiments. JT and EJD analyzed the data. JT and GJR drafted the manuscript. All authors contributed to editing and approving the final manuscript.

## Data and Code Accessibility

All analysis code and data referenced in this manuscript is available at via the Zenodo Digital Repository (Toxopeus et al., 2023), https://doi.org/10.5281/zenodo.10277669.

## Literature Cited

Andrewartha, H. G. 1952. Diapause in relation to the ecology of insects. Biological Reviews 27:50–107.

Bradshaw, W. E., and L. P. Lounibos. 1977. Evolution of dormancy and its photoperiodic control in pitcher-plant mosquitoes. Evolution 31:546–567.

Buckley, L. B. 2022. Temperature-sensitive development shapes insect phenological responses to climate change. Current Opinion in Insect Science 52:100897.

Bush, G. L. 1969. Sympatric host race formation and speciation in frugivorous flies of the genus *Rhagoletis* (Diptera, Tephritidae). Evolution 23:237–251.

Calvert, M. B., M. M. Doellman, J. L. Feder, G. R. Hood, P. Meyers, S. P. Egan, T. H. Q. Powell, et al. 2022. Genomically correlated trait combinations and antagonistic selection contributing to counterintuitive genetic patterns of adaptive diapause divergence in *Rhagoletis* flies. Journal of Evolutionary Biology 35:146–163.

Cayton, H. L., N. Haddad, K. Gross, S. E. Diamond, and L. Ries. 2015. Do growing degree days predict phenology across butterfly species? Ecology 96:1473–1479.

Chen, C., Q.-W. Xia, S. Fu, X.-F. Wu, and F.-S. Xue. 2014. Effect of photoperiod and temperature on the intensity of pupal diapause in the cotton bollworm, *Helicoverpa armigera* (Lepidoptera: Noctuidae). Bulletin of Entomological Research 104:12.

Chen, Y.-S., C. Chen, H.-M. He, Q.-W. Xia, and F.-S. Xue. 2013. Geographic variation in diapause induction and termination of the cotton bollworm, *Helicoverpa armigera* Hübner (Lepidoptera: Noctuidae). Journal of Insect Physiology 59:855–862.

Chino, H. 1958. Carbohydrate metabolism in the diapause egg of the silkworm, *Bombyx mori*—II: Conversion of glycogen into sorbitol and glycerol during diapause. Journal of Insect Physiology 2:1–12.

Dambroski, H. R., and J. L. Feder. 2007. Host plant and latitude-related diapause variation in *Rhagoletis pomonella*: a test for multifaceted life history adaptation on different stages of diapause development. Journal of Evolutionary Biology 20:2101–2112.

Damos, P., and M. Savopoulou-Soultani. 2012. Temperature-driven models for insect development and vital thermal requirements. Psyche: A Journal of Entomology 2012:1–13.

Dean, R. L., and P. J. Chapman. 1973. Bionomics of the apple maggot in eastern New York. Search Agriculture 3:1–64.

Denlinger, D. L. 2002. Regulation of diapause. Annual Review of Entomology 47:93–122.

Denlinger, D. L. 2022. Insect Diapause. Cambridge University Press, Cambridge.

Dingle, H., and T. A. Mousseau. 1994. Geographic variation in embryonic development time and stage of diapause in a grasshopper. Oecologia 97:179–185.

Dowle, E. J., T. H. Q. Powell, M. M. Doellman, P. J. Meyers, M. B. Calvert, K. K. O. Walden, H. M. Robertson, et al. 2020. Genome-wide variation and transcriptional changes in diverse developmental processes underlie the rapid evolution of seasonal adaptation. Proceedings of the National Academy of Sciences 117:23960–23969.

Dupuis, J. R., B. A. Mori, and F. A. Sperling. 2016. *Trogus* parasitoids of *Papilio* butterflies undergo extended diapause in western Canada (Hymenoptera, Ichneumonidae). Journal of Hymenoptera Research 50:179.

Emerson, K. J., W. E. Bradshaw, and C. M. Holzapfel. 2010. Microarrays reveal early transcriptional events during the termination of larval diapause in natural populations of the mosquito, *Wyeomyia smithii*. PLoS One 5:e9574.

Feder, J. L., T. A. Hunt, and L. Bush. 1993. The effects of climate, host plant phenology and host fidelity on the genetics of apple and hawthorn infesting races of *Rhagoletis pomonella*. Entomologia Experimentalis et Applicata 69:117–135.

Feder, J. L., U. Stolz, K. M. Lewis, W. Perry, J. B. Roethele, and A. Rogers. 1997. The effects of winter length on the genetics of apple and hawthorn races of *Rhagoletis pomonella* (Diptera: Tephritidae). Evolution 51:1862–1876.

Green, D. A., and M. R. Kronforst. 2019. Monarch butterflies use an environmentally sensitive, internal timer to control overwintering dynamics. Molecular Ecology 28:3642–3655.

Hahn, D. A., and D. L. Denlinger. 2011. Energetics of insect diapause. Annual Review of Entomology 56:103–121.

Hand, S. C., D. L. Denlinger, J. E. Podrabsky, and R. Roy. 2016. Mechanisms of animal diapause: recent developments from nematodes, crustaceans, insects, and fish. American Journal of Physiology 310:R1193–R1211.

Hanski, I. 1988. Four kinds of extra long diapause in insects: a review of theory and observations. Annales Zoologici Fennici 25:37–53.

Higaki, M., and M. Toyama. 2012. Evidence for reversible change in intensity of prolonged diapause in the chestnut weevil *Curculio sikkimensis*. Journal of Insect Physiology 58:56–60.

Hilbert, D. W., J. A. Logan, and D. M. Swift. 1985. A unifying hypothesis of temperature effects on egg development and diapause of the migratory grasshopper, *Melanoplus sanguinipes* (Orthoptera: Acrididae). Journal of Theoretical Biology 112:827–838.

Hodek, I. 1996. Diapause development, diapause termination and the end of diapause. European Journal of Entomology 93:475–487.

Hodek, I. 2002. Controversial aspects of diapause development. European Journal of Entomology 99:163–173.

Irwin, J. T., V. A. Bennett, and R. E. Lee. 2001. Diapause development in frozen larvae of the goldenrod gall fly, Eurosta solidaginis Fitch (Diptera: Tephritidae). Journal of Comparative Physiology B 171:181–188.

Jenkins, J., J. A. Powell, J. A. Logan, and B. J. Bentz. 2001. Low seasonal temperatures promote life cycle synchronization. Bulletin of Mathematical Biology 63:573–595.

Kankare, M., D. J. Parker, M. Merisalo, T. S. Salminen, and A. Hoikkala. 2016. Transcriptional differences between diapausing and non-diapausing *D. montana* females reared under the same photoperiod and temperature. PLoS One 11:e0161852.

Kim, M., R. M. Robich, J. P. Rinehart, and D. L. Denlinger. 2006. Upregulation of two actin genes and redistribution of actin during diapause and cold stress in the northern house mosquito, *Culex pipiens*. Journal of Insect Physiology 52:1226–1233.

King, A. M., and T. H. MacRae. 2012. The small heat shock protein p26 aids development of encysting *Artemia* embryos, prevents spontaneous diapause termination and protects against stress. PLoS One 7:e43723.

Kingsolver, J. G., G. J. Ragland, and J. G. Shlichta. 2004. Quantitative genetics of continuous reaction norms: thermal sensitivity of caterpillar growth rates. Evolution 58:1521– 1529.

Kingsolver, J. G., and H. A. Woods. 2016. Beyond thermal performance curves: modeling time-dependent effects of thermal stress on ectotherm growth rates. The American Naturalist 187:283–294.

Kipyatkov, V. E., and E. B. Lopatina. 2010. Intraspecific variation of thermal reaction norms for development in insects: new approaches and prospects. Entomological Review 90:163–184.

Koštál, V. 2006. Eco-physiological phases of insect diapause. Journal of Insect Physiology 52:113–127.

Koštál, V., T. Štětina, R. Poupardin, J. Korbelová, and A. W. Bruce. 2017. Conceptual framework of the eco-physiological phases of insect diapause development justified by transcriptomic profiling. Proceedings of the National Academy of Sciences 114:8532–8537.

Koštál, V., M. Tollarová, and D. Doležel. 2008. Dynamism in physiology and gene transcription during reproductive diapause in a heteropteran bug, *Pyrrhocoris apterus*. Journal of Insect Physiology 54:77–88.

Kutcherov, D. 2020. Stagewise resolution of temperature-dependent embryonic and postembryonic development in the cowpea seed beetle *Callosobruchus maculatus* (F.). BMC Ecology 20:50.

Lehmann, P., P. Pruisscher, V. Koštál, M. Moos, P. Šimek, S. Nylin, R. Agren, et al. 2018. Metabolome dynamics of diapause in the butterfly *Pieris napi*: distinguishing maintenance, termination and post-diapause phases. Journal of Experimental Biology 221:jeb169508.

Lehmann, P., W. Van Der Bijl, S. Nylin, C. W. Wheat, and K. Gotthard. 2017. Timing of diapause termination in relation to variation in winter climate. Physiological Entomology 42:232–238.

Lindestad, O., L. von Schmalensee, P. Lehmann, and K. Gotthard. 2020. Variation in butterfly diapause duration in relation to voltinism suggests adaptation to autumn warmth, not winter cold. Functional Ecology 34:1029–1040.

Lopatina, E. B., D. A. Kutcherov, and S. V. Balashov. 2014. The influence of diet on the duration and thermal sensitivity of development in the linden bug *Pyrrhocoris apterus* L. (Heteroptera: Pyrrhocoridae). Physiological Entomology 39:208–216.

Lyons-Sobaski, S., and S. H. Berlocher. 2009. Life history phenology differences between southern and northern populations of the apple maggot fly, *Rhagoletis pomonella*. Entomologia Experimentalis et Applicata 130:149–159.

Marshall, K. E., K. Gotthard, and C. M. Williams. 2020. Evolutionary impacts of winter climate change on insects. Current Opinion in Insect Science 41:54–62.

Masaki, S. 1980. Summer diapause. Annual Review of Entomology 25:1–25.

Masaki, S. 2002. Ecophysiological consequences of variability in diapause intensity. European Journal of Entomology 99:143–154.

Meyers, P. J., T. H. Q. Powell, K. K. O. Walden, A. J. Schieferecke, J. L. Feder, D. A. Hahn, H. M. Robertson, et al. 2016. Divergence of the diapause transcriptome in apple maggot flies: winter regulation and post-winter transcriptional repression. Journal of Experimental Biology 219:2613–2622.

Moraiti, C. A., C. T. Nakas, and N. T. Papadopoulos. 2014. Diapause termination of *Rhagoletis cerasi* pupae is regulated by local adaptation and phenotypic plasticity: escape in time through bet-hedging strategies. Journal of Evolutionary Biology 27:43–54.

Nielsen, M. E., P. Lehmann, and K. Gotthard. 2022. Longer and warmer prewinter periods reduce post-winter fitness in a diapausing insect. Functional Ecology 36:1151–1162.

Nomura, M., and Y. Ishikawa. 2000. Biphasic effect of low temperature on completion of winter diapause in the onion maggot, *Delia antiqua*. Journal of Insect Physiology 46:373–377.

Powell, J. A., and B. J. Bentz. 2009. Connecting phenological predictions with population growth rates for mountain pine beetle, an outbreak insect. Landscape Ecology 24:657–672.

Powell, T. H. Q., A. Nguyen, Q. Xia, J. L. Feder, G. J. Ragland, and D. A. Hahn. 2020. A rapidly evolved shift in life-history timing during ecological speciation is driven by the transition between developmental phases. Journal of Evolutionary Biology 33:1371–1386.

Prokopy, R. J. 1968. Influence of photoperiod, temperature, and food on initiation of diapause in the apple maggot. The Canadian Entomologist 100:318–329.

Pruisscher, P., P. Lehmann, S. Nylin, K. Gotthard, and C. W. Wheat. 2022. Extensive transcriptomic profiling of pupal diapause in a butterfly reveals a dynamic phenotype. Molecular Ecology 31:1269–1280.

R Core Team. 2022. R: A Language and Environment for Statistical Computing. R Foundation for Statistical Computing, Vienna, Austria.

Ragland, G. J., D. L. Denlinger, and D. A. Hahn. 2010. Mechanisms of suspended animation are revealed by transcript profiling of diapause in the flesh fly. Proceedings of the National Academy of Sciences 107:14909–14914.

Ragland, G. J., J. Fuller, J. L. Feder, and D. A. Hahn. 2009. Biphasic metabolic rate trajectory of pupal diapause termination and post-diapause development in a tephritid fly. Journal of Insect Physiology 55:344–350.

Ragland, G. J., and E. Keep. 2017. Comparative transcriptomics support evolutionary convergence of diapause responses across Insecta. Physiological Entomology 42:246–256.

Ragland, G. J., S. B. Sim, S. Goudarzi, J. L. Feder, and D. A. Hahn. 2012. Environmental interactions during host race formation: host fruit environment moderates a seasonal shift in phenology in host races of *Rhagoletis pomonella*. Functional Ecology, 26:921–931.

Réaumur, R. A., 1735. Observations du thermomètre faites pendant l’année MDCCXXXV comparées a celles qui ont été faites sous la ligne a l’Isle-de-France, a Alger et en quelques-unes de nos Isles de l’Amérique. Mémoires de l’Académie Royale des Sciences, 1735:545–576.

Rebaudo, F., and V.-B. Rabhi. 2018. Modeling temperature-dependent development rate and phenology in insects: review of major developments, challenges, and future directions. Entomologia Experimentalis et Applicata 166:607–617.

Régnière, J., J. Powell, B. Bentz, and V. Nealis. 2012. Effects of temperature on development, survival and reproduction of insects: experimental design, data analysis and modeling. Journal of Insect Physiology 58:634–647.

Reissig, W. H., J. Barnard, R. W. Weires, E. H. Glass, and R. W. Dean. 1979. Prediction of apple maggot fly emergence from thermal unit accumulation. Environmental Entomology 8:51–54.

Reynolds, J. A., and S. C. Hand. 2009. Embryonic diapause highlighted by differential expression of mRNAs for ecdysteroidogenesis, transcription and lipid sparing in the cricket *Allonemobius socius*. Journal of Experimental Biology 212:2075–2084.

Rinehart, J. P., A. Li, G. D. Yocum, R. M. Robich, S. A. Hayward, and D. L. Denlinger. 2007. Up-regulation of heat shock proteins is essential for cold survival during insect diapause. Proceedings of the National Academy of Sciences 104:11130–11137.

Roberts, K. T., N. E. Rank, E. P. Dahlhoff, J. H. Stillman, and C. M. Williams. 2021. Snow modulates winter energy use and cold exposure across an elevation gradient in a montane ectotherm. Global Change Biology 27:6103–6116.

Robich, R. M., and D. L. Denlinger. 2005. Diapause in the mosquito *Culex pipiens* evokes a metabolic switch from blood feeding to sugar gluttony. Proceedings of the National Academy of Sciences 102:15912–15917.

Roff, D., 1980. Optimizing development time in a seasonal environment: the ‘ups and downs’ of clinal variation. Oecologia 45:202–208.

Roff, D.A., 1996. The evolution of threshold traits in animals. The Quarterly Review of Biology 71:3–35.

Rull, J., E. Tadeo, R. Lasa, and M. Aluja. 2016. The effect of winter length on survival and duration of dormancy of four sympatric species of *Rhagoletis* exploiting plants with different fruiting phenology. Bulletin of Entomological Eesearch 106:818–826.

Salminen, T. S., L. Vesala, A. Laiho, M. Merisalo, A. Hoikkala, and M. Kankare. 2015. Seasonal gene expression kinetics between diapause phases in *Drosophila virilis* group species and overwintering differences between diapausing and non-diapausing females. Scientific Reports 5:11197.

Schmidt, P. S., L. Matzkin, M. Ippolito, and W. F. Eanes. 2005. Geographic variation in diapause incidence, life-history traits, and climatic adaptation in *Drosophila melanogaster*. Evolution 59:1721–1732.

Scranton, K., and P. Amarasekare. 2017. Predicting phenological shifts in a changing climate. Proceedings of the National Academy of Sciences 114:13212–13217.

Sgolastra, F., J. Bosch, R. Molowny-Horas, S. Maini, and W. P. Kemp. 2010. Effect of temperature regime on diapause intensity in an adult-wintering Hymenopteran with obligate diapause. Journal of Insect Physiology 56:185–194.

Shi, P., F. Ge, Y. Sun, and C. Chen. 2011. A simple model for describing the effect of temperature on insect developmental rate. Journal of Asia-Pacific Entomology 14:15–20.

Shingleton, A. W., G. C. Sisk, and D. L. Stern. 2003. Diapause in the pea aphid (*Acyrthosiphon pisum*) is a slowing but not a cessation of development. BMC Developmental Biology 3:7.

Sinclair, B. J. 2015. Linking energetics and overwintering in temperate insects. Journal of Thermal Biology 54:5–11.

Sinclair, B. J., K. E. Marshall, M. A. Sewell, D. L. Levesque, C. S. Willett, S. Slotsbo, Y. Dong, et al. 2016. Can we predict ectotherm responses to climate change using thermal performance curves and body temperatures? Ecology Letters 19:1372–1385.

Sinclair, B. J., C. M. Williams, and J. S. Terblanche. 2012. Variation in thermal performance among insect populations. Physiological and Biochemical Zoology 85:594–606.

Sridhar, V., and P. V. R. Reddy. 2013. Use of degree days and plant phenology: a reliable tool for predicting insect pest activity under climate change conditions. Pages 287–294 in H. C. P. Singh, N. K. S. Rao, and K. S. Shivashankar, eds. Climate-Resilient Horticulture: Adaptation and Mitigation Strategies. Springer, India.

Tauber, M. J., C. A. Tauber, and S. Masaki. 1986. Seasonal Adaptations of Insects. Oxford University Press, New York.

Tauber, M., and C. Tauber. 1976. Insect seasonality: diapause maintenance, termination, and postdiapause development. Annual Review of Entomology 21:81–107.

Taylor, F. 1981. Ecology and evolution of physiological time in insects. The American Naturalist 117:1–23.

Toxopeus, J., L. Gadey, L. Andaloori, M. Sanaei, and G. J. Ragland. 2021. Costs of averting or prematurely terminating diapause associated with slow decline of metabolic rates at low temperature. Comparative Biochemistry and Physiology A 255:110920.

Toxopeus, J., E. J. Dowle, L. Andaloori, and G. J. Ragland. 2023. Data and code from: Variation in thermal sensitivity of diapause development among individuals and over time predicts life history timing in a univoltine insect. American Naturalist, Zenodo Digital Repository, 10.5281/zenodo.10277669

Trudgill, D. L., A. Honek, D. Li, and N. M. van Straalen. 2005. Thermal time - concepts and utility. Annals of Applied Biology 146:1–14.

Urbanski, J. M., J. B. Benoit, M. R. Michaud, D. L. Denlinger, and P. Armbruster. 2010. The molecular physiology of increased egg desiccation resistance during diapause in the invasive mosquito, Aedes albopictus. Proceedings of the Royal Society B 277:2683–2692.

van Straalen, N. M. 1983. Physiological time and time-invariance. Journal of Theoretical Biology 104:349–357.

Vinogradova, E. B. 1986. Geographical variation and ecological control of diapause in flies. Pages 35–47 *in* F. Taylor and R. Karban, eds. The Evolution of Insect Life Cycles. Springer US, New York.

Williams, C. M., H. A. Henry, and B. J. Sinclair. 2015. Cold truths: how winter drives responses of terrestrial organisms to climate change. Biological Reviews 90:214–35.

Wilsterman, K., M. A. Ballinger, and C. M. Williams. 2021. A unifying, ecophysiological framework for animal dormancy. Functional Ecology 35:11–31.

Xiao, H., S. Wu, C. Chen, and F. Xue. 2013. Optimal low temperature and chilling period for both summer and winter diapause development in *Pieris melete*: based on a similar mechanism. PLoS One 8:e56404.

Yurk, B. P., and J. A. Powell. 2010. Modeling the effects of developmental variation on insect phenology. Bulletin of Mathematical Biology 72:1334–1360.

Zhao, M., C. Peng, W. Xiang, W. Deng, D. Tian, X. Zhou, G. Yu, H. He, and Z. Zhao. 2013. Plant phenological modeling and its application in global climate change research: overview and future challenges. Environmental Reviews 21:1–14.

Zhu, D., Y. Yang, and Z. Liu. 2009. Reversible change in embryonic diapause intensity by mild temperature in the Chinese rice grasshopper, *Oxya chinensis*. Entomologia Experimentalis et Applicata 133:1–8.

Blanckenhorn, W. U. 1997. Altitudinal life history variation in the dung flies *Scathophaga stercoraria* and *Sepsis cynipsea*. Oecologia 109:342–352.

Cherrill, A. J., and M. Begon. 1989. Timing of life cycles in a seasonal environment: the temperature-dependence of embryogenesis and diapause in a grasshopper (*Chorthippus brunneus* Thunberg). Oecologia 78:237–241.

Vargas, R. I., W. A. Walsh, E. B. Jang, J. W. Armstrong, and D. T. Kanehisa. 1996. Survival and development of immature stages of four Hawaiian fruit flies (Diptera: Tephritidae) reared at five constant temperatures. Annals of the Entomological Society of America 89:64–69.

