## Supplementary Information for "Variation in thermal sensitivity of diapause development among individuals and over time predicts life history timing in a univoltine insect"

**Appendix A: Simulation Model Building**

*Relationships between diapause development rates, temperature, and development duration*

Development rates (*R*) are often expressed as proportion development per day ($d^{-1}$), and the time (d) to complete development (*Y*) is simply equal to the inverse of that development rate ($Y = 1/R$) (Kipyatkov and Lopatina 2010; Shi et al. 2011; Damos and Savopoulou-Soultani 2012). The classic method to determine thermal sensitivity of a development rate is to rear groups of individuals at different temperatures and calculate development rate at each temperature as the inverse of development duration (Kipyatkov and Lopatina 2010). However, we can assess the relative breadth of thermal sensitivity of diapause development, as well as shifts in thermal sensitivity over time by using the following experimental design. Groups of diapausing individuals are exposed to a cold temperature (e.g., 4°C) for different lengths of time (*X*), followed by exposure to a warm temperature (e.g., 21°C) until evidence of post-diapause morphogenesis is observed. These temperatures must be selected such that post-diapause morphogenesis can only occur at the warm temperature (rate of morphogenesis at the cold temperature = 0). We can represent *R* as the rate of diapause development at the cold temperature, *r* as the rate of diapause development at the warm temperature, and $r_g$ as the rate of post-diapause morphogenesis at the warm temperature. Total time to complete development following a range of chilling durations can then be modelled in several ways, depending on the thermal sensitivity of diapause development.

<u>A single diapause timer: narrow vs. broad thermal sensitivity</u>

If diapause development rate has a narrow thermal sensitivity such that diapause only progresses at low temperatures (e.g., Fig. 2B), the rate of diapause development at the warm temperature is zero, i.e., $r = 0$. Therefore, any cold exposure duration that is less than the time required to complete diapause at 4°C ($1/R$) will result in a total development duration ($Y$) of zero because diapause is unable to complete (and no post-diapause morphogenesis is observed) at the warm temperature:

$$Y = 0 \text{ when } X < \frac{1}{R} \quad \text{(Eq'n S1.1)}$$

If an individual is kept at the cold temperature for a sufficient time to complete diapause ($1/R$), post-diapause morphogenesis will resume when transferred to the permissive warm temperature, and total development duration will be:

$$Y = X + \frac{1}{r_g} \text{ when } X \geq \frac{1}{R} \quad \text{(Eq'n S1.2)}$$

where $1/r_g$ is the chronological time required to complete post-diapause morphogenesis at the warm temperature. If an individual is chilled for longer than is needed to complete diapause (> $1/R$), that individual will enter post-diapause quiescence, and any additional chilling will increase total development duration with a slope of 1. That is, any additional chilling will prolong development time by the duration of the additional chilling.

Conversely, we can model a broad thermal sensitivity of diapause development rate, i.e., diapause can progress at both cold and warm temperatures (e.g., Fig. 2C and D). In this case, if the individual is never chilled:

$$Y = \frac{1}{r} + \frac{1}{r_g} \text{ when } X = 0 \quad \text{(Eq'n S2.1)}$$

where $1/r$ is the duration of diapause at the warm temperature. Conversely, if the individual is chilled for sufficient time to complete diapause development:

$$Y = X + \frac{1}{r_g} \text{ when } X \geq \frac{1}{R} \quad \text{(Eq'n S2.2)}$$

If an individual is chilled for longer than is needed to complete diapause, that individual will enter post-diapause quiescence, which will increase total development duration with a slope of 1. As described for Equation S1.2, any additional chilling will prolong development time by the duration of the additional chilling. If the individual is chilled at the cold temperature for a duration < $1/R$, some proportion of diapause development ($X \times R$) completes in the cold, and the remaining proportion of diapause development ($1 - X \times R$) completes during post-chill warming. The total duration of diapause development is therefore:

$$Y = X + \frac{1-(X \times \mathrm{R})}{r} + \frac{1}{r_g} \text{ when } 0 \leq X < \frac{1}{R} \quad \text{(Eq'n S2.3)}$$

which simplifies to a classic linear relationship ($y = mx + b$) between $X$ and $Y$:

$$Y = \frac{r - R}{r} X + \frac{1}{r} + \frac{1}{r_g} \text{ when } 0 \leq X < \frac{1}{R} \quad \text{(Eq'n S2.4)}$$

Using this equation, we see that the slope will be negative if chilling accelerates diapause development ($R > r$; Fig. 2C), and positive if warming accelerates diapause development ($r > R$; Fig. 2D).

Two diapause timers: incorporating a shift in thermal sensitivity during ontogeny

If thermal sensitivity of development can change during ontogeny, we must introduce additional diapause development rates. $R_e$ and $r_e$ are the rates of diapause development at cold and warm temperatures (respectively) early in diapause development, and $R_l$ and $r_l$ are the rates of diapause development at cold and warm temperatures (respectively) late in diapause development. The relationship between chill time ($X$) and total development duration ($Y$) in this more complicated scenario can be broken down into early and late diapause sections.

First, let's consider a narrow thermal sensitivity for our early diapause timer, such that early diapause only progresses at low temperatures. Any cold exposure duration that is less than the chronological time required to complete early diapause at 4°C ($1/R_e$) will result in a total development duration ($Y$) of zero because early diapause is unable to complete at the warm temperature (preventing any further development), similar to Equation S1.1:

$$Y = 0 \text{ when } X < \frac{1}{R_e} \quad \text{(Eq'n S3.1)}$$

Conversely, we can consider a broad thermal sensitivity for our early diapause timer. For chill durations that are insufficient to complete early diapause, some proportion of early diapause development ($X \times R_e$) completes in the cold, and the remaining proportion of early diapause development ($1 - X \times R_e$) completes during post-chill warming. In this case, the total duration of diapause is:

$$Y = X + \frac{1-(X \times R_e)}{r_e} + \frac{1}{r_l} + \frac{1}{r_g} \text{ when } X < \frac{1}{R_e} \quad \text{(Eq'n S3.2)}$$

where $X$ is the duration of partial early diapause development in the cold, $(1 - X \times R_e)/r_e$ is the time to complete the rest of early diapause at the warm temperature, and $1/r_l$ is the duration of late diapause development at the warm temperature. This simplifies to a classic linear relationship ($y = mx + b$) between $X$ and $Y$:

$$Y = \frac{r_e - R_e}{r_e} X + \frac{1}{r_e} + \frac{1}{r_l} + \frac{1}{r_g} \text{ when } X < \frac{1}{R_e} \quad \text{(Eq'n S3.3)}$$

Using this equation, we see that the slope will be negative if chilling accelerates diapause development ($R_e > r_e$; Fig. 2C), and positive if warming accelerates diapause development ($r_e > R_e$; Fig. 2D).

For the late diapause timer, we can consider the same two scenarios as for our early diapause timer. If the late diapause timer has a narrow thermal sensitivity (diapause only progresses at low temperatures), any cold exposure duration that is less than the time required to complete early *and* late diapause at 4°C ($1/R_e + 1/R_l$) will result in a total development duration ($Y$) of zero because diapause is unable to complete at the warm temperature, similar to Equation S1.1:

$$Y = 0 \text{ when } X < \frac{1}{R_e} + \frac{1}{R_l} \quad \text{(Eq'n S3.4)}$$

This would result in a development duration pattern that is indistinguishable from a single diapause timer that also has a cold-optimized narrow thermal sensitivity. However, if the late diapause timer has a broad thermal sensitivity, chilling that is sufficiently long for completion of early diapause but too short for completion of late diapause will result in the following:

$$Y = X + \frac{1-([X-\frac{1}{R_e}] \times R_l)}{r_l} + \frac{1}{r_g} \text{ when } \frac{1}{R_e} \leq X < \frac{1}{R_e} + \frac{1}{R_l} \quad \text{(Eq'n S3.5)}$$

where $X$ is the duration of early diapause and partial late diapause development in the cold, and $(1 - [X - 1/R_e] \times R_l)/r_l$ is the time to complete the rest of late diapause at the warm temperature. This simplifies to a classic linear relationship ($y = mx + b$) between $X$ and $Y$:

$$Y = \frac{r_l - R_l}{r_l} X + \frac{R_l + R_e}{R_e r_l} + \frac{1}{r_g} \text{ when } \frac{1}{R_e} \leq X < \frac{1}{R_e} + \frac{1}{R_l} \quad \text{(Eq'n S3.6)}$$

Using this equation, we see that the slope will be negative if chilling accelerates late diapause development ($R_l > r_l$; Fig. 2C), and positive if warming accelerates diapause development ($r_l > R_l$; Fig. 2D).

When individuals are chilled for a sufficient duration to complete diapause or for longer, total development duration will be expressed similar to Equations S1.2 and S2.2:

$$Y = X + \frac{1}{r_g} \text{ when } X \geq \frac{1}{R_e} + \frac{1}{R_l} \quad \text{(Eq'n S3.7)}$$

This relationship holds as long as both diapause processes can complete at 4°C, and does not depend on whether the early and late diapause timers have a narrow or broad thermal sensitivity. More complicated models are also possible (e.g., three or more diapause timers, each with their own thermal sensitivity), but we were able to simulate a population closely predicting empirical observations in *R. pomonella* using a maximum of two diapause timers per diapause phenotype.

*Simulation of Rhagoletis pomonella diapause development*

We calculated total development duration (since time zero) for each simulated individual (and whether that individual would complete development) under a range of chill durations using the assumptions in Table S1 and the equations in the main text. The rationale behind these equations and assumptions (as well as tests of those assumptions) is explained in the subsections below. Each individual in the population was simulated with its own diapause development rates at 4°C and 21°C, and these rates were sampled from simulated distributions of diapause development rates. The population distributions (mean and variation) of these simulated diapause rates are summarized in Table S2. We used different diapause timers and distributions of diapause development rates for weak diapause (WD) and chill-dependent diapause (CD) individuals, treating WD and CD as discrete phenotypes. We did consider simulations in which all individuals were sampled from the same distributions of diapause development rates (rather than having discrete WD and CD phenotypes), but doing so did not produce patterns that were consistent with the empirical data (see Alternative thermal sensitivity patterns below).

**Table S1. Assumptions for development of simulated weak diapause and chill-dependent diapause pupae chilled at 4°C at time zero (10 d post-pupariation) for different durations, followed by warming at 21°C.**

| | **Weak diapause** | **Chill-dependent diapause** |
|---|---|---|
| Thermal sensitivity | Diapause progresses across a wide range of temperatures and is accelerated by warming. | Early diapause only progresses well at low temperatures, while late diapause progresses across a wide range of temperatures and is accelerated by warming. |
| Variation in development rates | Diapause development rates at 4°C ($R$) and 21°C ($r$) have an inverse-Gaussian distribution. | The rate of early diapause development at 4°C ($R_e$) has an inverse-Gaussian distribution.<br>Early diapause development cannot progress at 21°C ($r_e$ = 0, or very close to 0). |
| | The ratio of the rate at 4°C to the rate at 21°C does not vary among individuals. | The rates of late diapause development at 4 °C ($R_l$) and 21 °C ($r_l$) are correlated with the rate of early diapause development. |
| | The rate of post-diapause morphogenesis at 21°C ($r_g$) is invariant. | The rate of post-diapause morphogenesis at 21°C ($r_g$) is invariant. |
| Mortality due to chilling | Pupae die if chilled for $X_z$ days. | Pupae die if chilled for $X_z$ days. |

**Table S2. Model parameters of a simulated population of 150 weak diapause and 850 chill-dependent diapause pupae.** Development rates were modelled with inverse Gaussian distributions with mean μ and shape λ (Fig. S4). Corresponding mean ($\bar{x}$) and standard deviation ($s$) of durations to complete diapause or a component of diapause (early and late) at a single temperature (4°C and 21°C, respectively) are shown in the same row. The effect of increasing or decreasing these rates can be seen in Fig. S7.

| | Development rates ($d^{-1}$) | | Development durations (d) | |
|---|---|---|---|---|
| | Parameter | Values | Parameter | Values |
| **Weak diapausers (*N* = 150)** | | | | |
| *At 21 °C* | $r$ | $\mu = 0.030,\ \lambda = 0.25$ | $\frac{1}{r}$ | $\bar{x} = 35.9, s = 11.9$ |
| *At 4 °C* | $R$ | $\mu = 0.017, \lambda = 0.1$ | $\frac{1}{R}$ | $\bar{x} = 65.7, s = 21.7$ |
| **Chill-dependent diapausers (*N* = 850)** | | | | |
| *At 21 °C* | $r_e$ | $\mu = 0$ | $\frac{1}{r_e}$ | 0 (no development) |
| | $r_l$ | $\mu = 0.014,\ \lambda = 10$ | $\frac{1}{r_l}$ | $\bar{x} = 71.0, s = 2.7$ |
| *At 4 °C* | $R_e$ | $\mu = 0.0155,\ \lambda = 0.3$ | $\frac{1}{R_e}$ | $\bar{x} = 68.2, s = 16.4$ |
| | $R_l$ | $\mu = 0.0035, \lambda = 3$ | $\frac{1}{R_l}$ | $\bar{x} = 285.6, s = 10.6$ |

Thermal sensitivity, mortality and equations for simulated weak diapause

We simulated our **weak diapause (WD)** individuals with a single diapause timer with a broad thermal sensitivity of diapause development (Table S1). We chose a broad thermal sensitivity for the following reasons. 1) Our empirical data (and that of others; Dambroski and Feder 2007; Calvert et al. 2022) shows that WD *R. pomonella* can eclose rapidly (35 – 65 days post-pupariation) with no chilling, suggesting diapause can complete at warm temperatures. 2) The proportion of pupae that eclosed with little to no chilling (0 – 4 weeks) was constant at approximately 15% (Fig. 3C), and approximately 15 % of our population consisted of WD pupae (Fig. 3B), suggesting that all flies that eclosed with 0 – 4 weeks chilling were WD individuals. 3) During these first 4 weeks of chilling, mean total development duration increased with chilling, but post-chill development duration decreased (yellow + gray in Fig. 4), suggesting diapause could progress at 4°C as well as 21°C. Further, the positive slope of mean development duration vs. chill duration during these first 4 weeks of chilling (Fig. 4) suggested that WD pupal diapause was accelerated by warm temperatures (cf. Fig. 2D). Note that two or more diapause timers, each with a broad thermal sensitivity, would also be consistent with the empirical observations associated with WD pupae. However, a single diapause timer was the most parsimonious option and was sufficient to produce a good fit to empirical data (see Results).

To model total development duration in our simulated WD pupae, we used Equation S2.4 (above), and Equations S2.5 – S2.6 (below) [Equations 1.1 – 1.3 in the main text]. We know that WD pupae are killed by moderate chilling (Toxopeus et al. 2021), so we also incorporated a mortality term into our final model. Equation S2.5 is a slightly modified Equation S2.2. that incorporates this mortality:

$Y = X + \frac{1}{r_g}$ when $\frac{1}{R} \leq X < X_z$ (Eq'n S2.5)

Where $X_z$ is the chill duration that causes mortality in that WD individual. Chill durations longer than $X_z$ resulted in a total development duration of zero because no eclosion was observed:

$Y = 0$ when $X \geq X_z$ (Eq'n S2.6)

See Fig. S8 for the impacts of removing WD mortality from our model.

Thermal sensitivity and equations for simulated chill-dependent diapause

We simulated our **chill-dependent diapause (CD)** individuals with two diapause timers: an early diapause timer with a narrow thermal sensitivity and a late diapause timer with a broad thermal sensitivity of diapause development (Table S1). Our reasoning for this was as follows. 1) For moderate chill times (6 – 14 weeks), as chill time increased proportion eclosion increased in an approximately sigmoidal pattern (Fig. 3C). We moved forward with the likely assumption that this trend in proportion eclosion was driven by CD pupae, as WD pupae seemed to exhibit a proportion eclosion pattern that did not depend on chilling. Therefore, CD pupae seemed to require chilling (e.g., due to a narrow thermal sensitivity of diapause development) early in development. 2) We saw no evidence for post-diapause quiescence in the chilling treatments we used in our study; even our longest chill treatment (29 weeks) resulted in post-chill development duration that took longer than the mean time to complete morphogenesis at 21°C (Fig. 4). Therefore, although chilling was required early in diapause development, there seemed to be a later diapause process with a broad thermal sensitivity – one that could complete at either 4°C or 21°C. Further, the positive slope of mean development duration vs. chill duration following prolonged chilling (17 – 29 weeks) (Fig. 4) suggested that this late diapause process in CD pupae was accelerated by warm temperatures (cf. Fig. 2D). Additional diapause timers (e.g., three or more) are possible, but two single diapause timers was the most parsimonious option, and was sufficient to produce a good fit to empirical data while allowing insight into ontogenetic shifts in the thermal sensitivity of diapause development (see Results).

To model total development duration in our simulated CD pupae, we used Equations S3.1 and S3.6 (above), and Equations S3.8 – S3.9 (below) [Equations 2.1 – 2.4 in the main text]. We saw increased mortality after prolonged chilling (Fig. S1) that was likely due to death of CD pupae (because WD pupae likely die after moderate chilling). Equation S3.8 is a slightly modified Equation S3.7. that incorporates mortality:

$$Y = X + \frac{1}{r_g} \text{ when } \frac{1}{R_e} + \frac{1}{R_l} \leq X < X_z \qquad \text{(Eq'n 3.8)}$$

where $X_z$ is the chill duration that causes CD pupa mortality. Chill durations longer than $X_z$ resulted in a total development duration of zero because no eclosion was observed:

$$Y = 0 \text{ when } X \geq X_z \qquad \text{(Eq'n 3.9)}$$

See Fig. S8 for the impacts of removing CD mortality from our models.

Variation in diapause development rates

Because development duration generally had a Gaussian distribution (Fig. S2), we simulated variation in diapause development rates ($r$, $R$, $R_e$, $r_l$, $R_l$) among individuals with inverse Gaussian distributions (Table S1). Normal distributions of diapause development rates did not substantially improve the simulation model fit to empirical data (Fig. S4).

We assumed that different rates of diapause development (at 4°C vs. 21°C, early vs. late diapause) were correlated within an individual (Table S1). This assumption is based on principles and examples from the morphogenesis development rate literature. For example, development durations across life history stages can be correlated within individuals such that some individuals consistently develop faster than others at multiple temperatures. This can be seen in yellow dung fly larvae vs. pupae (Blanckenhorn 1997) and grasshopper eggs early and late in embryogenesis (Cherrill and Begon 1989). See Figs. S5 and S6 for the minimal impact of this assumption about rate correlation within an individual on model fit.

We allowed for a change in the magnitude of interindividual variation across temperatures (e.g., variation in $R$ [4°C] and $r$ [21°C]), and across ontogeny (e.g., variation in $R_e$ [early diapause] and $R_l$ [late diapause] at 4°C). Variation in morphogenic development rates can change across temperature; for example several fruit fly species shown less variation in pupal development rate at high temperatures than low temperatures (Vargas et al. 1996). The morphogenic development rate/time literature contains several examples of changes in the variability of development rate across life stages, e.g., egg development duration is less variable than pupal development duration in Mountain Pine Beetle at a given temperature (Yurk and Powell 2010). See Figs. S5 and S6 for the substantial impact of these assumptions about variation in rates across temperature and ontogeny, respectively, on model fit.

We did not model interindividual variation in duration of post-diapause morphogenesis (Table S1) because this post-diapause morphogenesis is minimally variant among individuals of

*R. pomonella* at a given temperature, and variation in eclosion time is generally driven by variation in diapause duration in *R. pomonella* (Powell et al. 2020).

*Alternative thermal sensitivity patterns*

While the simulation model outlined in Tables S1 and S2 effectively captured the general trends in mean total development duration, CoV development duration, and proportion eclosion, we considered whether alternative approaches to thermal sensitivity patterns could improve model fit.

First, we considered whether CD pupae could be modelled with a rate of early diapause development at 21°C that was greater than zero ($r_e > 0$). This was intended to account for previous observations that a small proportion of CD *R. pomonella* can eclose without chilling, although this typically takes more than 150 d (Dambroski and Feder 2007). When we calculated simulation summary metrics for our $r_e > 0$ model, the fit to empirical data was very poor (Fig. S9A vs. B). However, in our empirical data experiment, we did not track eclosion indefinitely, so we did not count any pupae that might have eclosed after a prolonged time post-chill. If we incorporated a 'cutoff' of 115 days into our $r_e > 0$ model – i.e., if we did not 'count' any simulated eclosion that occurred after 115 days post-chill – the fit of this model (Fig. S9C) was similar but not a dramatic improvement relative to our base model ($r_e = 0$; Fig. S9A).

Second, we considered whether we could model all pupae from a single set of diapause timers for early and late diapause. To do this, we hypothesized ways in which thermal sensitivity (thermal performance curve) of diapause development rate could vary continuously among individuals, such that some required chilling to complete diapause while others did not. For example, all individuals in a population might have the same thermal maximum temperature, but variation in breadth of thermal sensitivity (Fig. S10 A-B). In both hypothetical examples, it is possible to have a small number of individuals with a non-zero early diapause development rate at 21°C ($r_e > 0$; WD-like), and for all individuals to have a non-zero early diapause development rate at 4°C ($R_e > 0$; Figs. S10A, C). Thus, continuous variation in thermal sensitivity of early diapause development rate could produce discrete phenotypes (zero vs. non-zero rates of development at 21°C) if pupae are incubated at a temperature near the upper (or indeed lower) limits of the thermal sensitivity of most individuals.

Using the approach above (i.e., the development rate distributions in Fig. S10C), we were able to produce patterns of mean development duration with approximately the correct shape, i.e., a positive slope with a discontinuity associated with moderate chilling, but a poor fit for short to no chilling durations (Fig. S10D). The new model had a poor CoV fit (Fig. S10E), which suggests the approach did not appropriately capture the true nature of interindividual variation in development rates and durations. Proportion eclosion patterns were approximately appropriate (Fig. S10F), as long as mean $R_e$ did not deviate substantially from the final model described in the main text. Refinement of this alternative model to fit empirical data may be possible, but is not a trivial task, and still requires binary categories (zero vs. non-zero development at 21°C in early diapause) even if those categories are produced from an originally continuous distributions of thermal sensitivities. Thus, we make our inferences based on the simpler approach of modelling discrete WD and CD phenotypes.

### Supplementary Figures

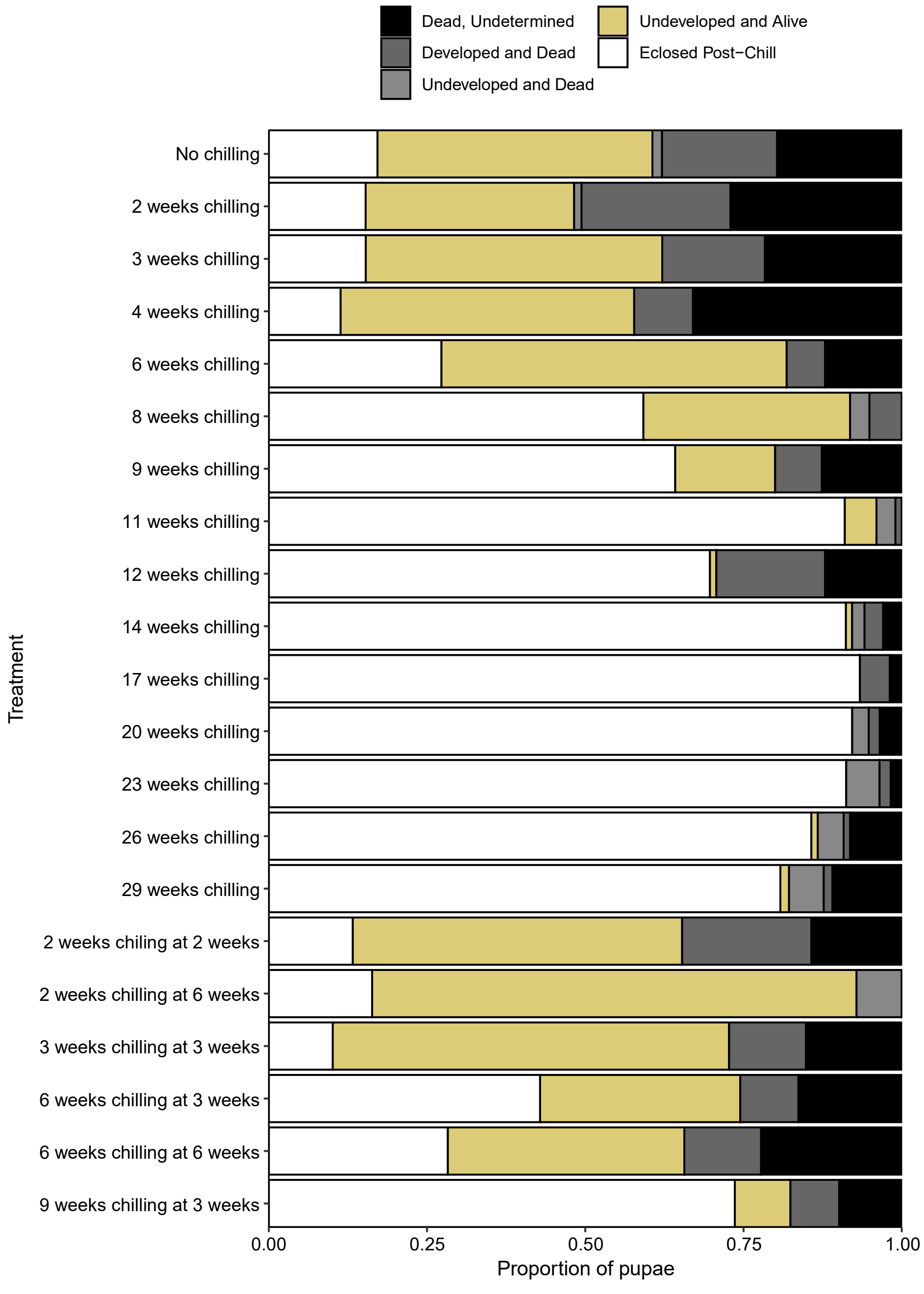


**Figure S1. Proportion of pupae that eclosed or remained uneclosed in each treatment condition.** Groups of pupae were exposed to the 4°C chilling conditions (e.g., 2 weeks) at the indicated time post-pupariation; if no age is indicated, the pupae were transferred to chilling at time zero (10 d post-pupariation). Pupae that eclosed are subdivided into those that eclosed (post-chill, except for the "No chilling" treatment) or did not eclose. Pupae that did not eclose are classified as undeveloped (still in diapause) and alive or dead, developed (completed diapause) and dead, and those that died but could not be developmentally staged (e.g., due to fungal growth or substantial decomposition).

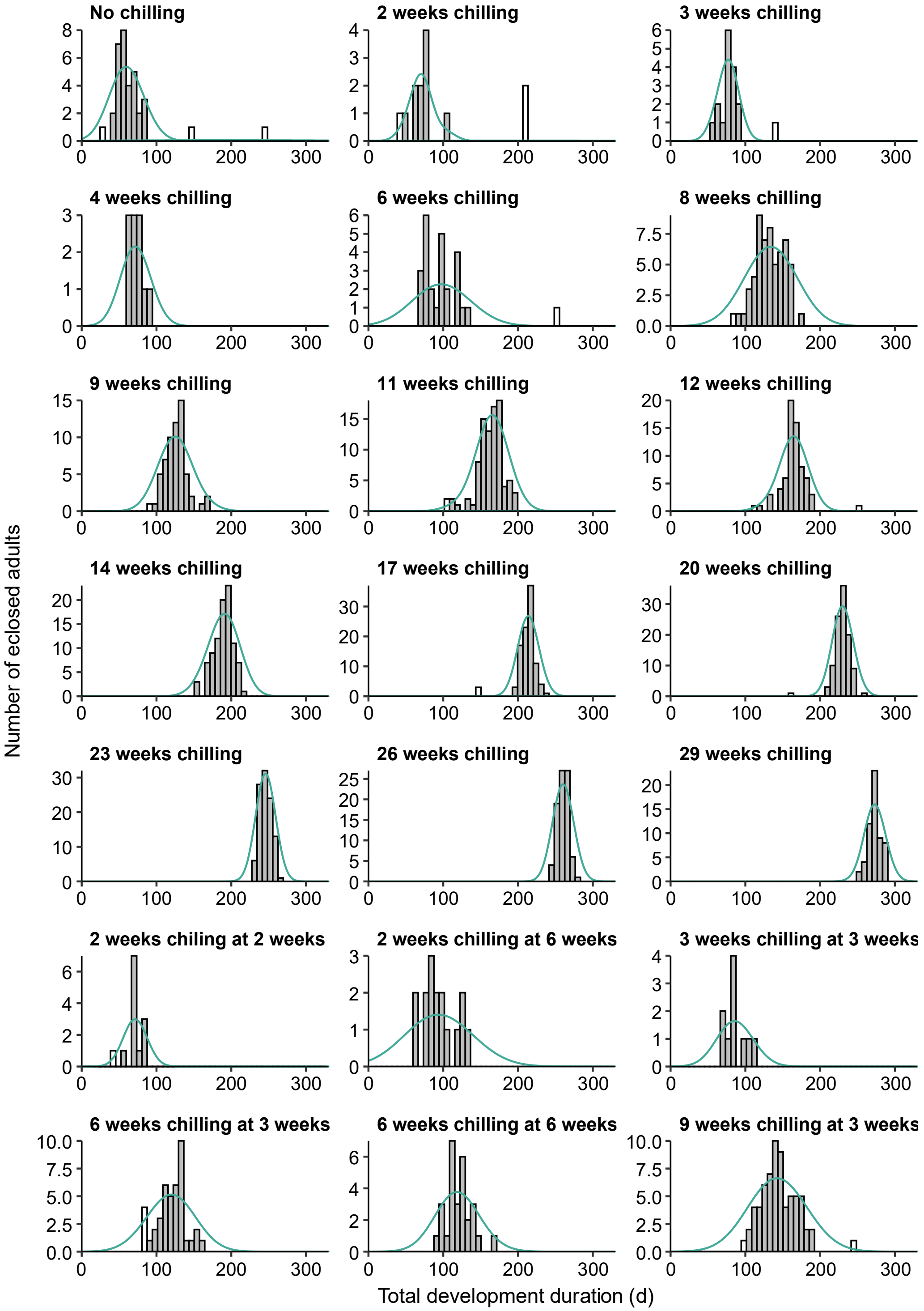


**Figure S2. Distributions of total development duration for each treatment condition.** Groups of pupae were exposed to the 4°C chilling conditions (e.g., 2 weeks) at time zero or the indicated age. White bars represent individuals who were excluded from further analyses because they were non-diapause pupae (eclosed very early) or had outlier eclosion times (eclosed very late). A normal distribution function (teal) is superimposed over each dataset.

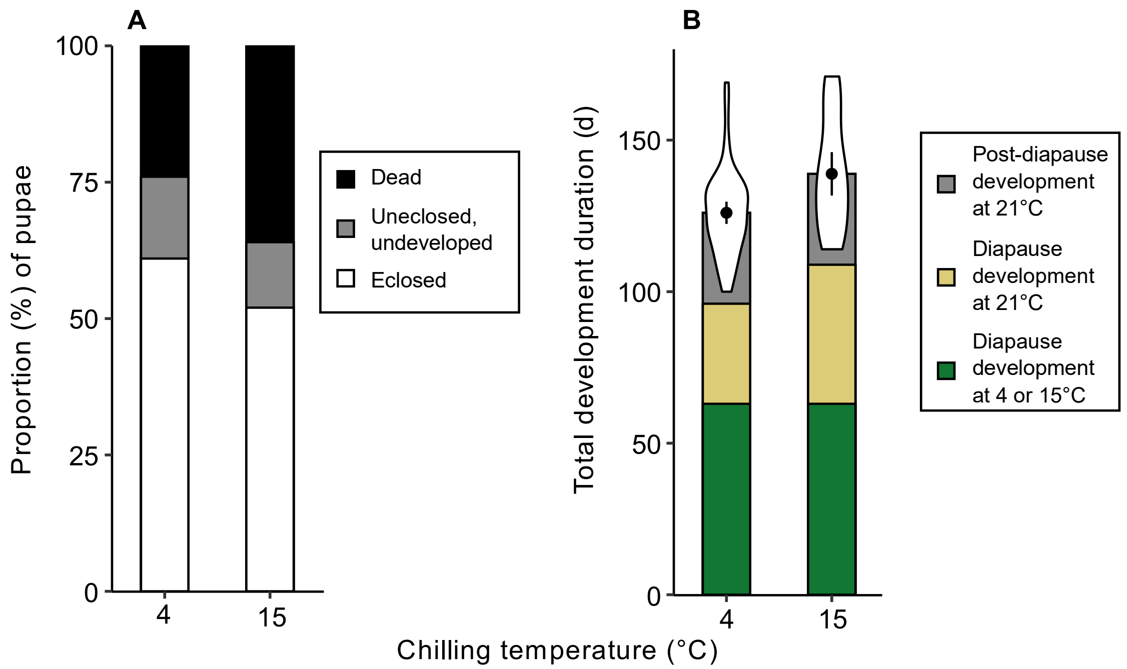


**Figure S3. The effect of chilling temperature on (A) the proportion of *R. pomonella* that complete diapause and post-diapause development, and (B) the duration of that development. (A)** Each bar represents the proportion of c. 100 pupae that eclosed as adults (white), failed to eclose but remained alive and in diapause (gray), or failed to eclose and died (black) following 9 weeks of chilling at 4°C or 15°C in darkness at time zero (10 d post-pupariation). **(B)** Each violin plot represents the distribution of development durations in pupae that eclosed as adults at 21°C following exposure to the same treatments as in **(A)**. Each point represents the mean ± 95% confidence intervals of total development duration. Bars below the violin plots show the duration of diapause development at 4°C or 15°C (green – dictated by chilling time), and estimated mean durations of diapause development (yellow) and post-diapause morphogenesis (gray) at 21°C.

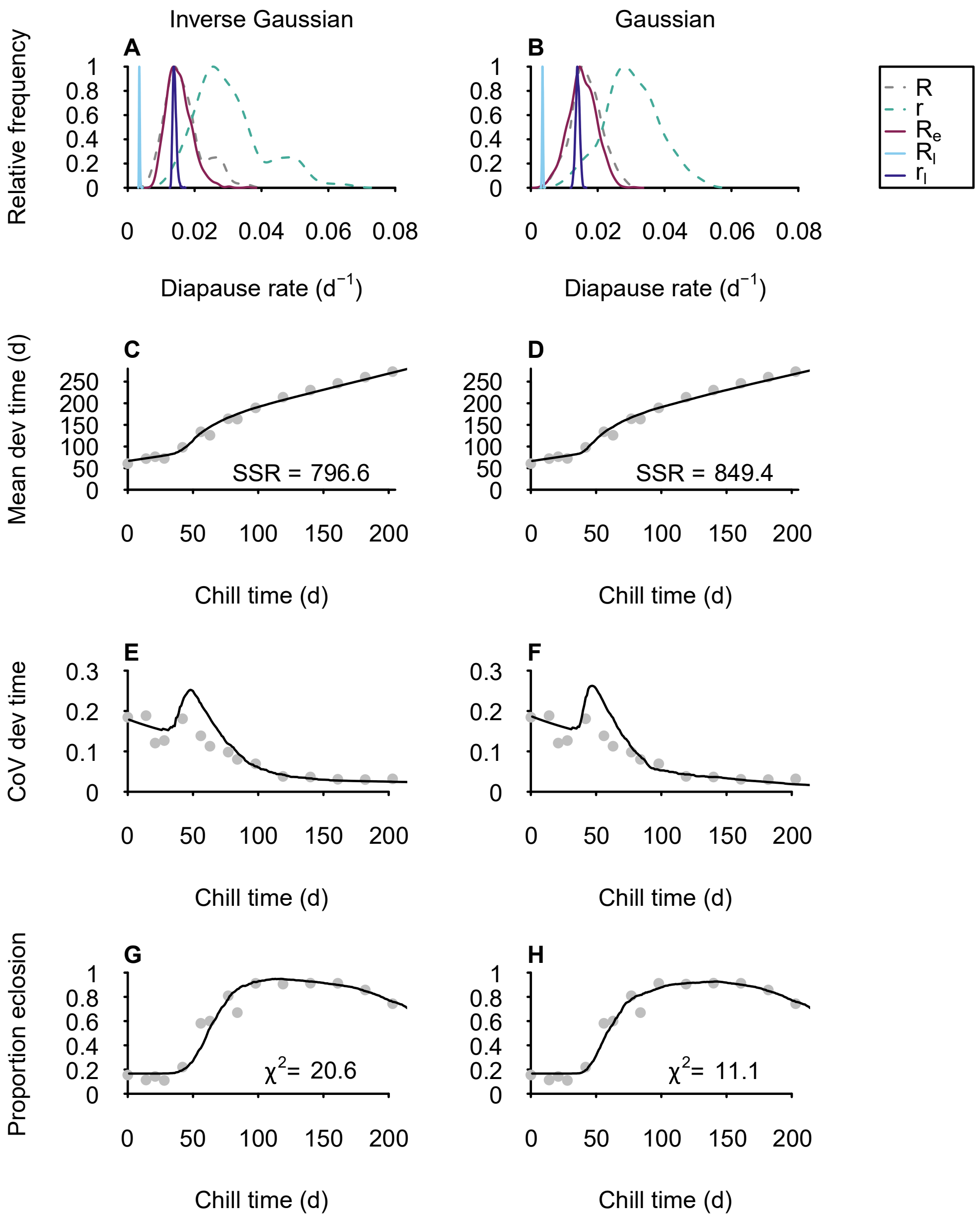


**Figure S4. Comparing simulated diapause development rates with (A) inverse Gaussian or (B) Gaussian distributions, and the effect of these distributions on (C – H) the simulation's fit to empirical data.** Diapause development rates include those for weak diapause at 4°C ($R$) and 21°C ($r$), early chill-dependent diapause at 4°C ($R_e$), and late chill-dependent diapause at 4°C ($R_l$) and 21°C ($r_l$). Empirical data are represented as gray points for the total development duration, coefficient of variation (CoV) of mean development duration, and proportion eclosion as a function of chill duration. The simulation calculations of these same variables are represented by the solid black lines, and were determined from the development rates in **A** (left column) or **B** (right column). SSR, sum of squared residuals; $\chi^2$, Chi squared.

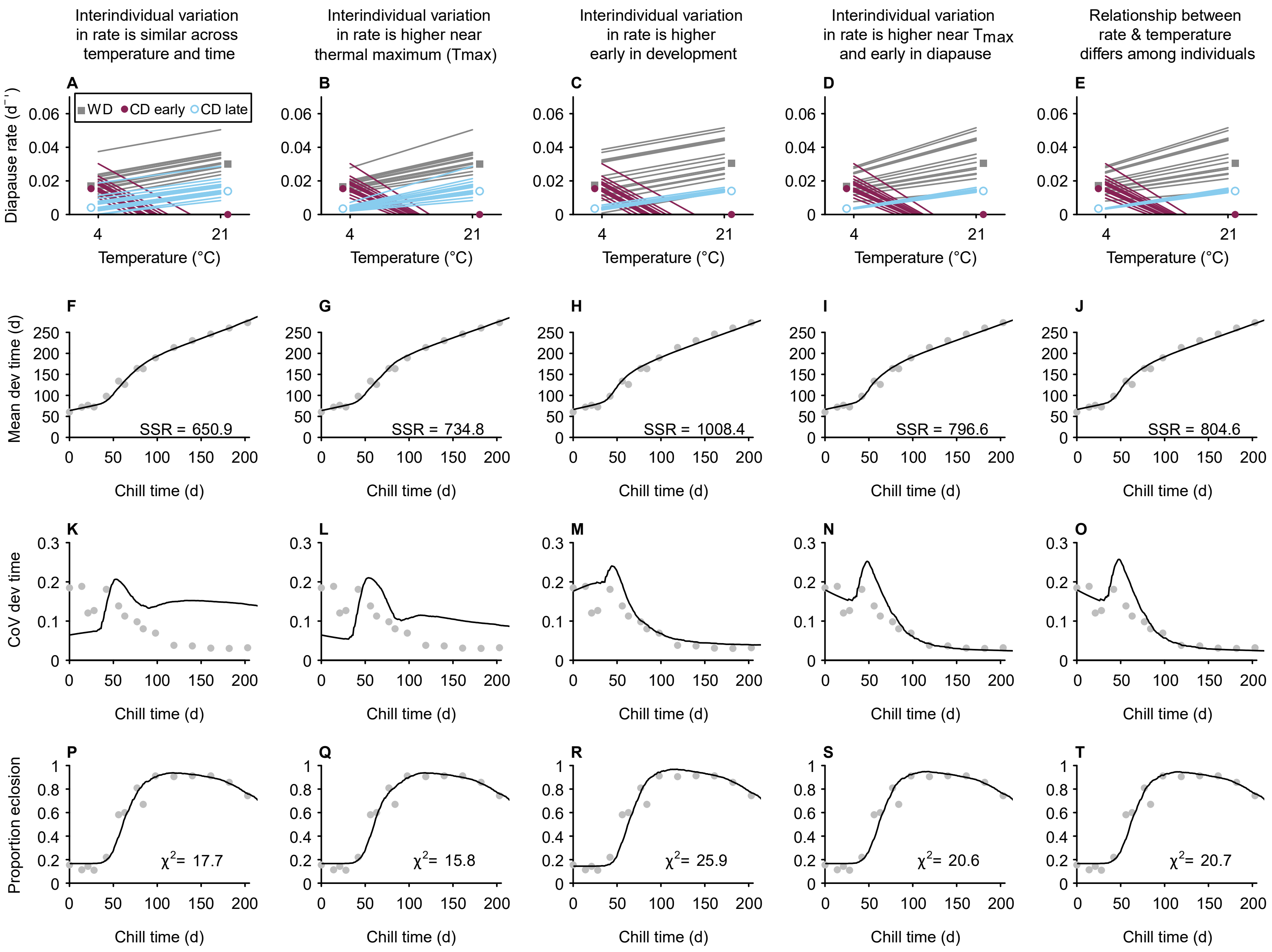

Interindividual variation in rate is similar across temperature and time
Interindividual variation in rate is higher near thermal maximum (Tmax)
Interindividual variation in rate is higher early in development
Interindividual variation in rate is higher near Tmax and early in diapause
Relationship between rate & temperature differs among individuals
A
B
C
D
E
WD
CD early
CD late
Diapause rate (d⁻¹)
Temperature (°C)
F
G
H
I
J
Mean dev time (d)
SSR = 650.9
SSR = 734.8
SSR = 1008.4
SSR = 796.6
SSR = 804.6
Chill time (d)
K
L
M
N
O
CoV dev time
P
Q
R
S
T
Proportion eclosion
χ2= 17.7
χ2= 15.8
χ2= 25.9
χ2= 20.6
χ2= 20.7

**Figure S5 (above). Comparing variation in (A – E) simulated diapause development rates *across temperature* for weak diapause (WD) and early or late chill-dependent diapause (CD) individuals, and the effect of that variation on (F – T) the simulation's fit to empirical data**. **(A – E)** Each line represents a simulated individual in WD or CD (early or late), and points represent the mean development rates of each group at the two temperatures. Only 15 simulated individuals per group are shown for clarity purposes. The simulations were run as described in the main text, and mean development rates in each diapause group at each temperature (4°C and 21°C) were held approximately constant, but the variation around the mean was defined in different ways for each "column" in this Figure. **(A)** Variation in development rate among individuals is similar at 4°C and 21°C, and in WD and CD pupae at all times during development. **(B)** Variation in development rate among individuals is highest near the thermal maximum ($T_{max}$) for a process, i.e., 4°C for CD early diapause and 21°C for WD and CD late diapause. We assumed that the thermal maximum of development was closest to which ever temperature had the higher development rate for a given process. **(C)** Variation in development rate among individuals is similar at 4°C and 21°C within a group, but early developmental processes (e.g., CD early diapause) show greater variation in developmental rate than later developmental processes (e.g., CD late diapause). **(D)** A combination of the two principles in **B** and **C**; the parameters for this simulation match the final model in the main text. **(E)** Although not readily apparent from the plot (due to scale), this simulation uses the same assumptions as **D**, but slopes of the lines within each group are allowed to differ. **(F – T)** Empirical data are represented as gray points for the mean development duration, coefficient of variation (CoV) of development duration, and proportion eclosion as a function of chill duration. The simulation calculations of these same metrics are represented by the solid black lines, and were determined from the development rates represented at the top of each column. SSR, sum of squared residuals; $\chi^2$, Chi squared.

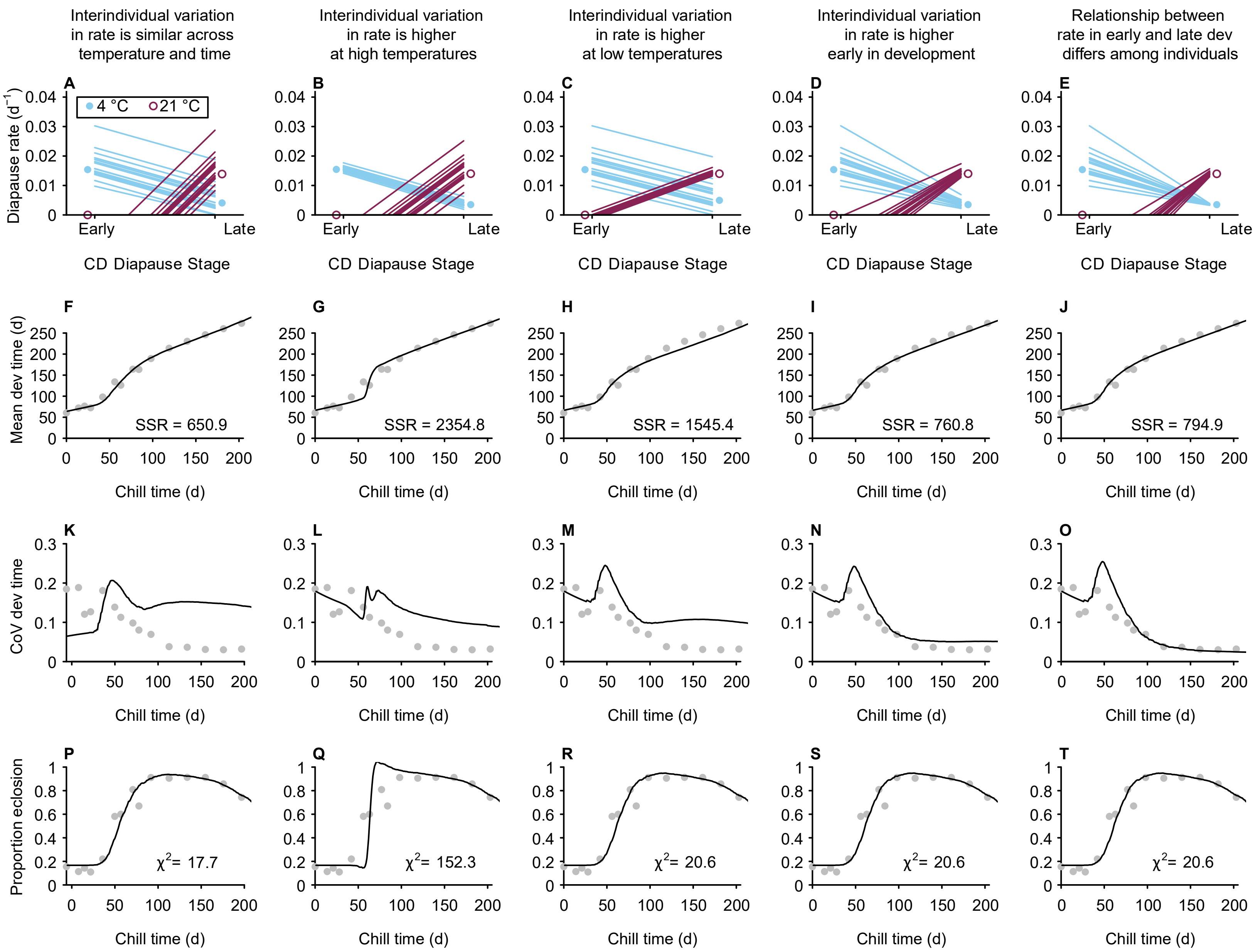
Interindividual variation in rate is similar across temperature and time
Interindividual variation in rate is higher at high temperatures
Interindividual variation in rate is higher at low temperatures
Interindividual variation in rate is higher early in development
Relationship between rate in early and late dev differs among individuals
A
B
C
D
E
4 °C
21 °C
Diapause rate (d⁻¹)
Early
Late
CD Diapause Stage
F
G
H
I
J
Mean dev time (d)
SSR = 650.9
SSR = 2354.8
SSR = 1545.4
SSR = 760.8
SSR = 794.9
Chill time (d)
K
L
M
N
O
CoV dev time
P
Q
R
S
T
Proportion eclosion
χ2= 17.7
χ2= 152.3
χ2= 20.6
χ2= 20.6
χ2= 20.6

**Figure S6 (above). Comparing variation in (A – E) simulated diapause development rates *across ontogeny* for chill-dependent diapause (CD) individuals, and the effect of that variation on (F – T) the simulation's fit to empirical data. (A – E)** Each line represents a simulated CD individual whose development rates are known during early and late diapause at 4°C and 21°C. Points represent the mean development rates at these two developmental stages. Only 15 simulated individuals per group are shown for clarity purposes. The simulations were run as described in the main text, and mean development rates in each group at each temperature (4°C and 21°C) were held approximately constant, but the variation around the mean was defined in different ways for each "column" in this Figure. Weak diapause (WD) individuals are not plotted but were included in each simulation using the exact same parameters as the final model described in the main text. **(A)** Interindividual variation in development rate is similar at 4°C and 21°C, and early and late diapause. **(B)** Variation in development rate among individuals is similar in early and late diapause at a given temperature, but variation at 21°C is higher than variation at 4°C. **(C)** As in **B**, except variation is higher at 4°C than 21°C. **(D)** Variation in development rate among individuals is higher in early diapause than late diapause. **(E)** Although not readily apparent from the plot (due to scale), this simulation uses the same assumptions as D, but slopes of the lines within each group are allowed to differ. **(F – T)** Empirical data are represented as gray points for the mean development duration, coefficient of variation (CoV) of development duration, and post-chill proportion eclosion as a function of chill duration. The simulation calculations of these same metrics are represented by the solid black lines. SSR, sum of squared residuals; $\chi^2$, Chi squared.

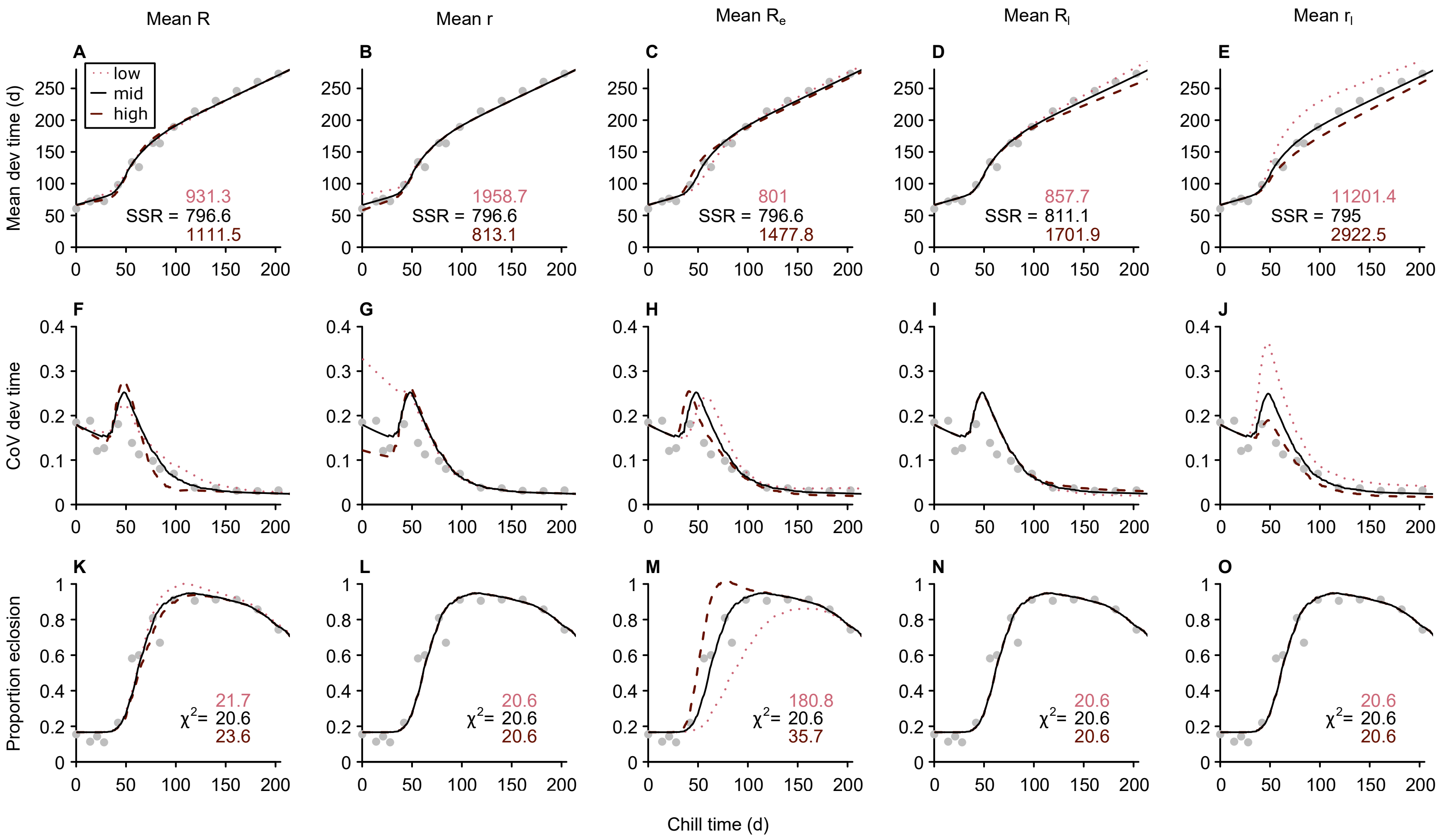


**Figure S7. Comparing the effect of increasing or decreasing the mean diapause development rate on the simulation's fit to empirical data.** Empirical data are represented as gray points for the **(A – E)** mean development duration, **(F – J)** coefficient of variation (CoV) of development duration, and **(K – O)** proportion eclosion as a function of chill duration. Within a "column," only the parameter indicated in the column title (*R*, *r*, $R_e$, $R_l$, $r_l$). was varied, while the other four rates were the same as the final model described in the main text. The effect of increasing ('low') or decreasing ('high') the mean simulated development rate relative to an intermediate rate ('mid') is shown. SSR, sum of squared residuals, and $\chi^2$, Chi squared, are colour-coded in the same way as the lines.

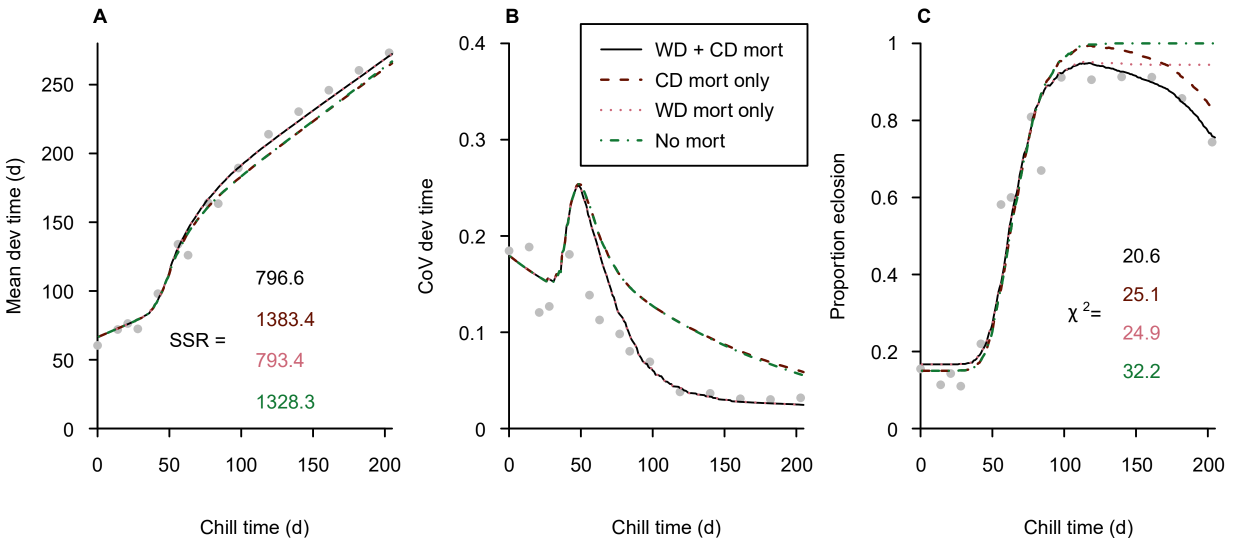


**Figure S8. Comparing the effect of mortality due to prolonged chilling in weak diapause (WD) and chill-dependent diapause (CD) individuals on the simulation's fit to empirical data.** Empirical data are represented as gray points for the (A) mean development duration, (B) coefficient of variation (CoV) of development duration, and (C) proportion eclosion as a function of chill duration. The final model in the main text (WD and CD mortality) is represented by the black line. Simulations that exclude WD mortality, CD mortality, or both are represented by the other lines. WD mortality only is indistinguishable from the final model in panels **A** and **B**. SSR, sum of squared residuals, and $\chi^2$, Chi squared, are colour-coded in the same way as the lines.

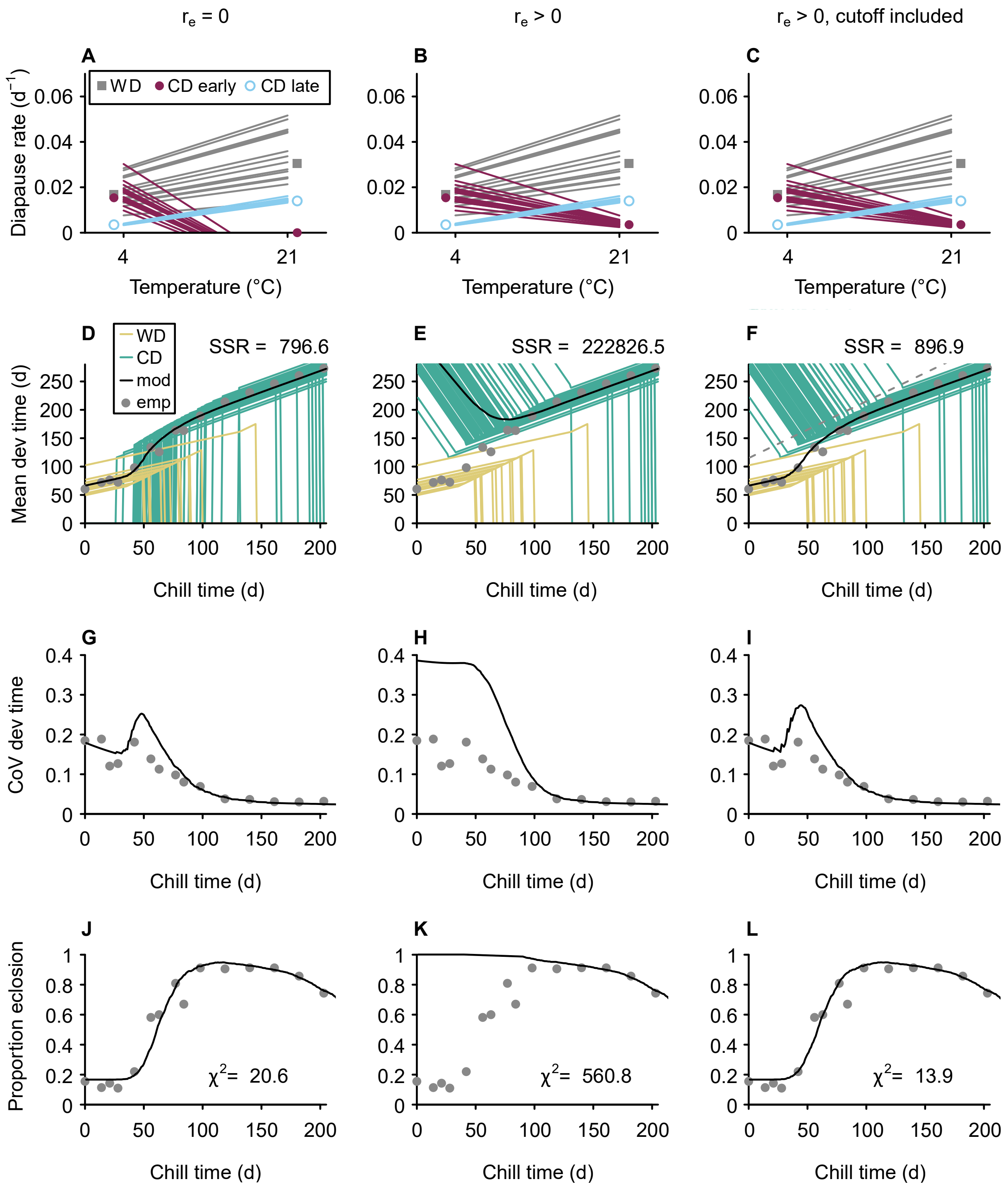

r_e = 0
r_e > 0
r_e > 0, cutoff included
A
B
C
WD
CD early
CD late
Diapause rate (d^-1)
Temperature (°C)
D
E
F
WD
CD
mod
emp
SSR = 796.6
SSR = 222826.5
SSR = 896.9
Mean dev time (d)
Chill time (d)
G
H
I
CoV dev time
J
K
L
Proportion eclosion
χ² = 20.6
χ² = 560.8
χ² = 13.9

**Figure S9 (above). The effect of (A – C) early diapause development rate at 21°C ($r_e$) in chill-dependent diapause (CD) individuals on (D – L) the simulation's fit to empirical data. (A – C)** Each line represents a simulated weak diapause (WD) or CD individual, and points represent the mean development rates of each group at the two temperatures. Only 15 simulated individuals per group are shown for clarity purposes. **(D – F)** Representative individuals are shown with yellow (WD) and green (CD) lines; vertical "drops" in the lines indicate mortality. **(D – L)** Gray points represent metrics calculated from empirical data: mean development duration, coefficient of variation (CoV) of development duration, and post-chill proportion eclosion. The simulation models (black lines) were run as described in the main text, and $r_e$ was varied as described in each "column." In the right-most column, simulated individuals that eclosed after a cutoff of 115 days post-chill (dashed line in **F**) were not included in the calculation of simulation summary metrics. SSR, sum of squared residuals; $\chi^2$, Chi squared. Additional interpretation of this figure is provided at the end of Appendix A.

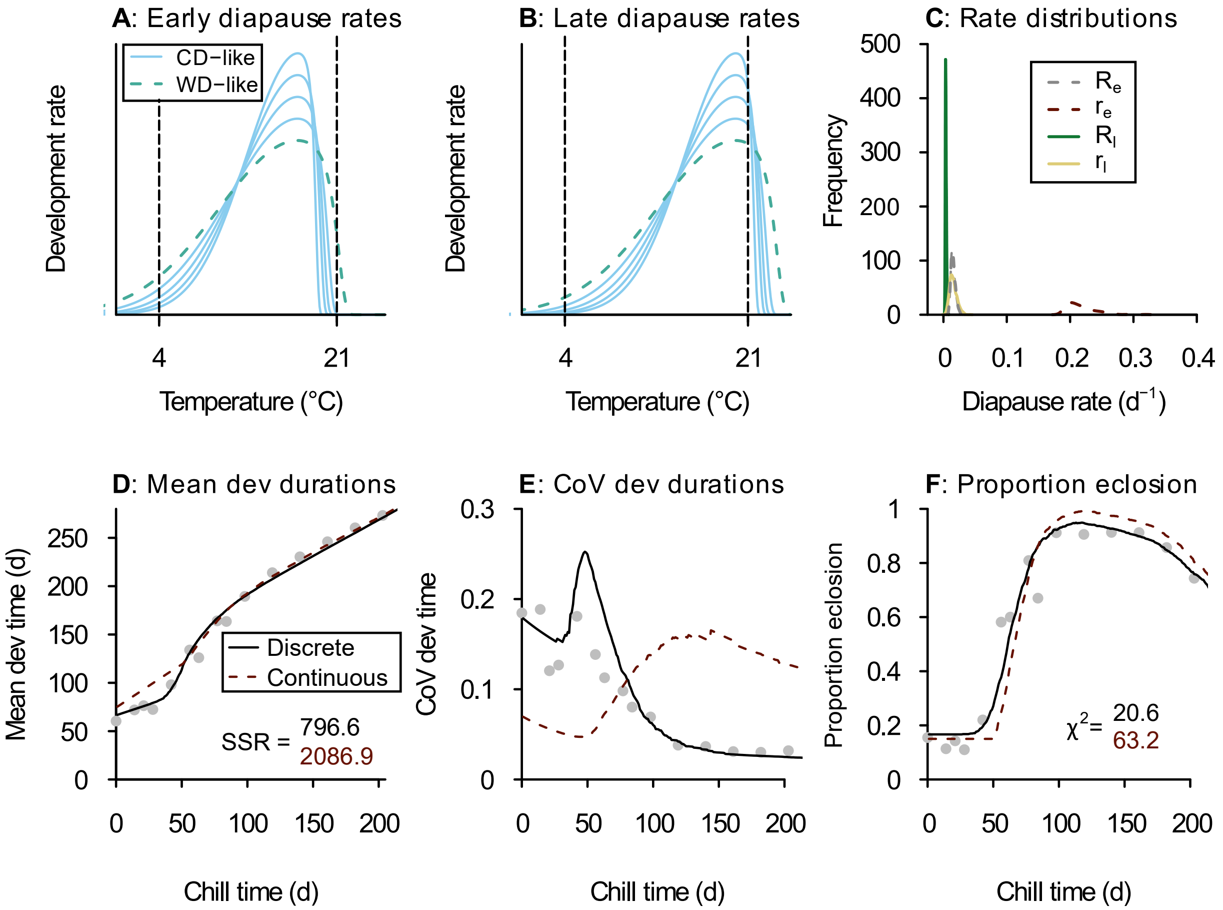


**Figure S10. Comparing the effect of (A – C) continuous variation in thermal sensitivity of diapause timers on (D – F) the simulation's fit to empirical data.** Hypothesized example patterns of interindividual variation of **(A)** early diapause and **(B)** late diapause development rate that could produce patterns similar to our empirical data. Each curve represents one individual; light blue solid line, similar to CD pupae; dashed teal green line, similar to WD pupae (non-zero development rate at 21°C). Dashed vertical lines included to highlight the range of development rates at the temperatures used in our study. **(C)** Distributions of diapause development rates used in our 'continuous' model. $R_e$, early diapause at 4°C; $r_e$, early diapause at 21°C; $R_l$, late diapause at 4°C; $r_l$, late diapause at 21°C. Note that only WD-like individuals have $r_e > 0$. **(D – F)** Gray points represent metrics calculated from empirical data: mean development duration, coefficient of variation (CoV) of development duration, and post-chill proportion eclosion. The 'continuous' model (dashed pink lines) based on rates in panel C is compared to the final simulation model (black lines) based on 'discrete' WD and CD phenotypes is described in the main text. SSR, sum of squared residuals, and $\chi^2$, Chi squared, are colour-coded in the same way as the lines. Additional interpretation of this figure is provided at the end of Appendix A.
